# Quantitative fluxomics of circulating metabolites

**DOI:** 10.1101/2020.03.02.973669

**Authors:** Sheng Hui, Alexis J. Cowan, Xianfeng Zeng, Lifeng Yang, Tara TeSlaa, Xiaoxuan Li, Caroline Bartman, Zhaoyue Zhang, Cholsoon Jang, Lin Wang, Wenyun Lu, Jennifer Rojas, Joseph Baur, Joshua D. Rabinowitz

## Abstract

Mammalian organs are nourished by nutrients carried by the blood circulation. These nutrients originate from diet and internal stores, and can undergo various interconversions before their eventual use as tissue fuel. Here we develop isotope tracing, mass spectrometry, and mathematical analysis methods to determine the direct sources of circulating nutrients, their interconversion rates, and eventual tissue-specific contributions to TCA cycle metabolism. Experiments with fifteen nutrient tracers enabled extensive accounting both for circulatory metabolic cycles and tissue TCA inputs, across fed and fasted mice on either high carbohydrate or ketogenic diet. We find that a majority of circulating carbon flux is carried by two major cycles: glucose-lactate and triglyceride-glycerol-fatty acid. Futile cycling through these pathways is prominent when dietary content of the associated nutrients is low, rendering internal metabolic activity robust to food choice. The presented *in vivo* flux quantification methods are broadly applicable to different physiological and disease states.

## INTRODUCTION

As animals, we get our nutrients from the food we eat. Tissues in our body, however, do not take their nutrients directly from food. Instead, they acquire nutrients from the blood circulation. Tissues must continuously burn circulating nutrients to extract energy for maintaining function, but animals do not consume food all the time. To resolve this mismatch, in the simplest scenario, animals store dietary nutrients in fed state and utilize the stored nutrients in fasted state (Figure 1A). Decades of research support this basic model, for example showing that after feeding insulin drives glucose storage as glycogen, while during fasting glucagon promotes release of glycogen to maintain blood glucose levels (Frayn, 2009; Newsholme and Leech, 2011).

**Figure 1.**
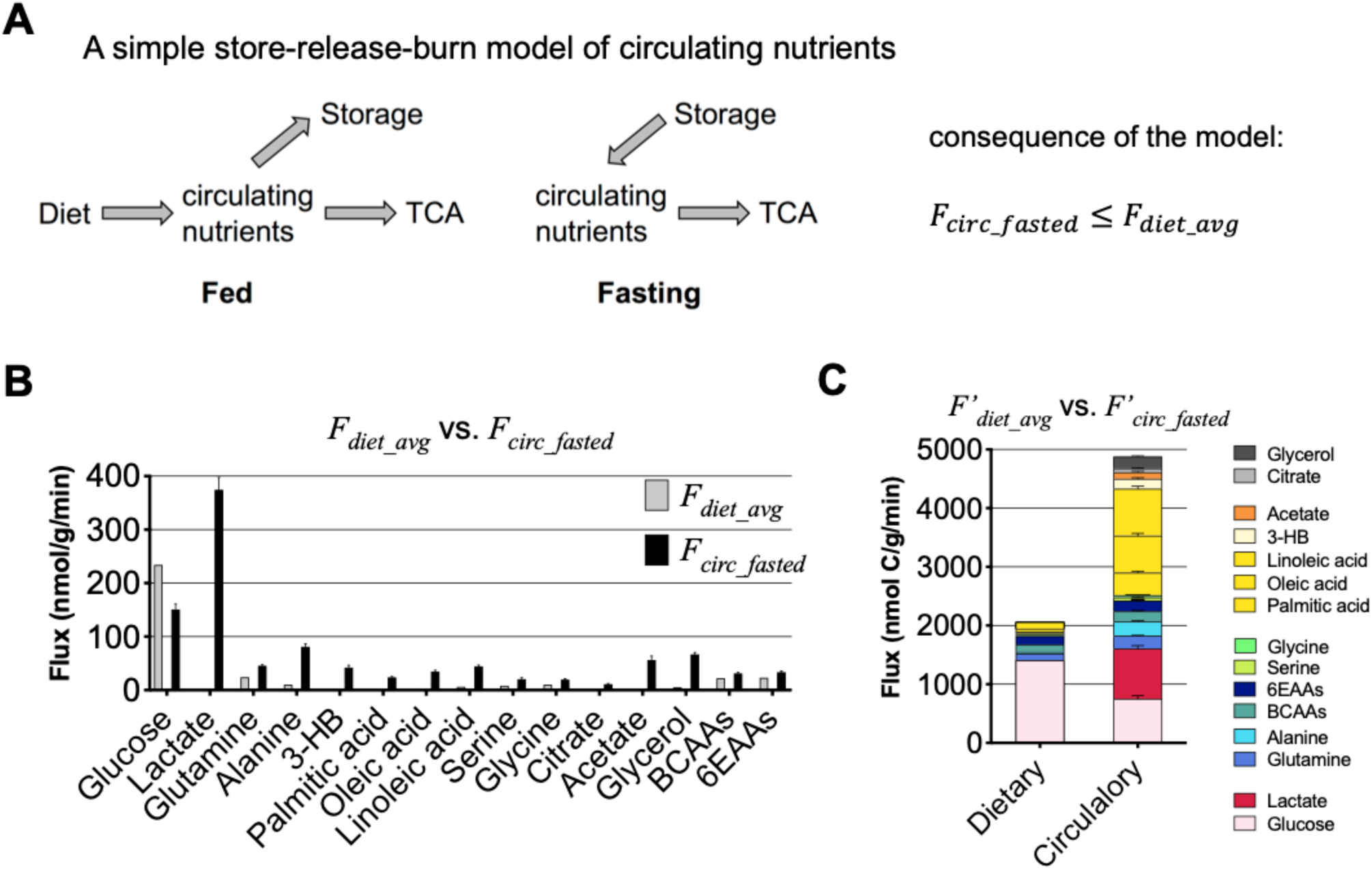
Circulatory turnover flux markedly exceeds dietary flux. Data are from 8-h fasted mice on carbohydrate diet. (A) Store-release-burn model. In fed state, circulating nutrients originate from the diet and go to either storage or tissue energy generation. In fasted state, circulating nutrients come from the storage and are consumed solely for tissue energy generation. For a nutrient that follows this model, in fasted state, its circulatory turnover flux is less than its average dietary intake flux averaged over a diurnal cycle. (B) Comparison between average dietary flux and fasted circulatory turnover flux for specific metabolites. See Table S1 for values of fluxes and N. (C) Comparison between the average dietary flux and fasted circulatory turnover flux, here shown as stacked bars. Different from (B), the flux units here are moles carbon, rather than moles molecules. See Table S1 for values.

At the same time, beyond simply being stored, released, and burned, circulatory metabolites are interconnected. For example, in addition to its direct production by the breakdown of hepatic glycogen, circulating glucose is also maintained during fasting by synthesis from lactate released by muscle. This glucose can then be used by muscle as fuel, creating a loop known as the Cori cycle (Frayn, 2009; Newsholme and Leech, 2011). Similarly, the Cahill cycle describes the interconversion between glucose and alanine (Felig, 1973). Another example is the glucose-glycerol interconnection in the triglyceride/fatty acid cycle, wherein the glycerol released by the breakdown of triglycerides can be used for glucose production and the glycerol backbone in the re-synthesized triglycerides can come from glucose (Reshef et al., 2003).

While there has been a great deal of progress in understanding these metabolic pathways and their regulation, it remains unclear whether metabolism largely follows the simple logic of nutrient storage, release, and burning, or whether interconversion of circulating metabolites is a major quantitative feature. Moreover, as animals make divergent food choices (e.g., a diet high in carbohydrate versus a diet high in fat), much of the impact on internal metabolic activities remains to be elucidated.

Here, we present systems-level methods for *in vivo* flux quantification and systematically determine the direct sources and interconversion fluxes of the most important circulating nutrients. By direct source, we mean the circulating precursor that is chemically converted into the metabolic product of interest without going through other circulating nutrients. We further trace circulating nutrients into the TCA cycle of diverse tissues, revealing tissue fuel preferences. By conducting these analyses for both fasted and fed mice on either standard carbohydrate-rich lab chow (carbohydrate diet, CD) or a ketogenic diet (KD), we provide a systems-level quantitative view of mammalian metabolic activity across divergent diets. The resulting data reveal fundamental features of metabolism. Specific examples include glycerol standing out among major circulating metabolites in not being a direct TCA fuel, carbohydrate (i.e., glucose and lactate) being the main brain fuel even on ketogenic diet, and heart being unique among organs in not burning circulating amino acids. More generally, we find that tissues have strong nutrient preferences that are maintained across feeding and fasting, and that major circulating nutrient cycles persist across physiological states and diets.

## RESULTS

### Comparison of Dietary Flux and Circulatory Flux

Consider the simple store-release-burn model shown in Figure 1a. In this model, tissues are fed by dietary nutrients (e.g. glucose, amino acids), either directly or after their storage (e.g. in glycogen, protein). Define *F*_*diet*_ as dietary intake rate of a particular nutrient (units of nmol/min/g, dividing the intake rate by the mouse weight in grams); *F*_*circ*_ as the rate of disappearance of the nutrient or other metabolite from the bloodstream due to tissue consumption, which at pseudo-steady-state is equivalent to its rate of appearance in the bloodstream from either diet or tissue excretion (see Methods) (Hui et al., 2017; Wolfe and Chinkes, 2005); and *F*_*burn*_ as the terminal clearance of the circulating nutrient, typically via burning into CO_2_ in the TCA cycle. Each of these parameters can be determined during the fed state, fasted state, or as an average rate throughout the day. Ultimately, over a diurnal cycle, organisms maintain metabolic pseudo-steady-state, and hence, in the store-release-burn model, *F*_*diet_avg*_ *F*_*burn_avg*_. Moreover, in the fasted state, nutrients taken up from the bloodstream are typically burned to generate energy rather than stored, and hence *F*_*circ_fasted*_ *F*_*burn_fasted*_. Finally, energy expenditure and thus nutrient burning is typically higher during feeding than fasting, and thus *F*_*circ_fasted*_ *F*_*burn_fasted*_ ≤ *F*_*burn_avg*_ *F*_*diet_avg*_. Thus, in the store-release-burn model, *F*_*circ_fasted*_ ≤ *F*_*diet_avg*_.

The features of the store-release-burn model can be compared to experimental flux data, to assess whether metabolism largely follows this logic, or operates in a fundamentally different way. In previous work, we identified the circulating metabolites in mice that carry an *F*_*circ*_ greater than 10% of the glucose *F*_*circ*_ (Hui et al., 2017). Here, in addition to these 13 top circulating metabolites, for comprehensiveness, we also include two other groups of metabolites, the three branched-chain amino acids (BCAAs) and six other essential amino acids (6EAAs), which were infused together in uniformly ^13^C-labeled form. Pyruvate was not included, as it is in rapid exchange with lactate and yet much less abundant in the bloodstream (Romijn et al., 1994).

Consistent with the store-release-burn model, glucose and the essential amino acids show a pattern where *F*_*circ_fasted*_ ≈ *F*_*diet_avg*_ (Figure 1B; Table S1). Most of circulatory metabolic flux, however, resides in nutrients for which *F*_*circ_fasted*_ >> *F*_*diet_avg*_: lactate, glutamine, fatty acids, glycerol, and acetate (Figure 1B; Table S1). Thus, metabolism is not primarily organized around storing, releasing, and burning dietary nutrients. Instead, interconversion between dietary nutrients and other circulating metabolic intermediates plays a large role.

### Alternative Measure of Fluxes

In contrast to fluxes measured in units of mole nutrient, to better reflect overall contributions of nutrients to carbon metabolism, in the rest of the paper we report flux in units of mole carbon, such that two lactate are equivalent to one glucose and one-third of a C18 fatty acid.

In addition, rather than reporting *F*_*circ*_, which uses mass spectrometry to measure any loss of *fully labeled infused tracer* (even if some of the labeled tracer atoms remain), we report 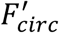 which we define as clearance of the *labeled tracer atoms* from the circulation (see Methods). 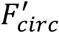 better aligns with classical radioactive tracer studies and facilitates certain downstream calculations (Wolfe and Chinkes, 2005). With the uniform units of nmol carbon/min/g for each nutrient, we can sum up the dietary fluxes and the circulatory turnover fluxes for an overall comparison (Figure 1C; Table S1). In the carbohydrate diet, diet flux is dominated by glucose from carbohydrate. Yet glucose, lactate, free fatty acids, glycerol, and amino acids all have high circulatory turnover flux. The substantial circulatory turnover flux of nutrients other than glucose reflects widespread interconversion between dietary nutrients, circulating nutrients, and internal nutrient stores like protein and triglycerides.

### Determining the Direct Sources of Circulating Metabolites

To determine the interconversion between dietary nutrients and circulating nutrients, we next aimed to quantify contributions of dietary nutrients to other circulating metabolites, and back again (in the case of metabolic cycles). To this end, for any circulating metabolite *X*, we wanted to figure out how much it contributes to any other circulating metabolite *Y*. The experimental approach is to infuse at a minimally perturbative rate uniformly ^13^C-labeled *X* and measure the pseudo-steady state labeling of *Y*. Specifically, upon infusion of *X*, we define the normalized labeling of *Y* (*L*_*Y*←*X*_) as fraction of labeled carbon atoms in serum *Y* relative to serum *X* (see Methods). For example, by infusing uniformly ^13^C-alanine and measuring labeling of circulating alanine and lactate, we can determine *L*_*lac*←*ala*_.

This measure of labeling, however, cannot be taken as the direct contribution from alanine to lactate, because the infused alanine may label lactate either directly or indirectly via other circulating metabolites (Figure. 2a). For example, alanine might first be taken into one tissue, converted glucose, excreted into the circulation, and then the circulating glucose might be converted into circulating lactate. Thus, the normalized labeling of lactate by alanine (*L*_*lac*←*ala*_) is the sum of the direct contribution from alanine to lactate (denoted as *f*_*lac*←*ala*_) and the indirect contribution through glucose, with the latter term equal to the direct contribution from glucose to lactate (*f*_*lac*←*glc*_) weighted by the normalized labeling of glucose by alanine (*L*_*glc*←*ala*_) (first equation in Figure 2A). Both direct contributions can be determined by separately infusing labeled alanine and labeled glucose and measuring the labeling of both serum lactate (*L*_*lac*←*glc*_) and alanine (*L*_*ala*←*glc*_) (second equation in Figure 2A). This allows the two equations to be solved for two direct contributions *f*_*lac*←*ala*_ and *f*_*lac*←*glc*_.

**Figure 2.**
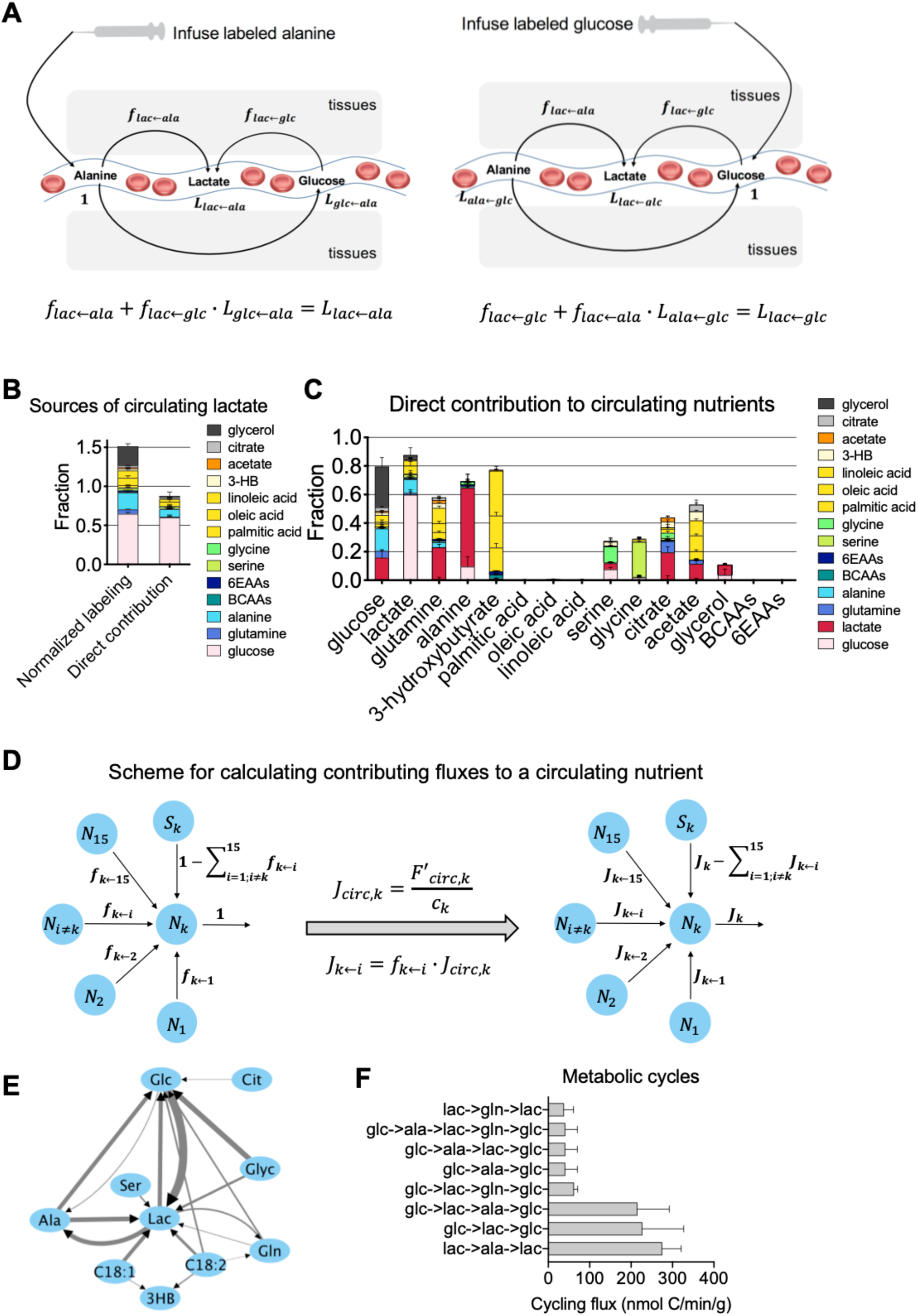
Comprehensive tracer studies enable determination of direct sources and interconversion fluxes of circulating metabolites. Data are for 8-h fasted mice on carbohydrate diet. (A) Example of calculating the direct contributions from two circulating metabolites (alanine, glucose) to a third metabolite (lactate). This requires infusion separately of alanine and of glucose. Each infusion experiment yields an isotope balance equation. Together, the two equations from the two infusions determine the two direct contributions. (B) Comparison between the normalized labeling of lactate (calculated separately from each infusion) and the direct contributions to lactate. (C) Direct contributions to each of 15 circulating metabolites from the other 14. See Table S2 for normalized labeling data and Table S3 for the direct contribution values. (D) Scheme for calculating absolute contributing fluxes from the relative direct contributions. (E) Circulating nutrient interconversion fluxes. Edge widths are proportional to log-transformed flux values. Shown are fluxes > 50 nmol C/min/g. (F) Metabolic cycles with flux > 50 nmol C/min/g.

This scheme can be extended to calculate the direct contribution from any circulating metabolite to any other circulating metabolite. To determine fully the inter-conversion between *n* circulating metabolites, we infuse each of them individually and measure their labeling in each infusion experiment. The resulting data form an *n* × *n* matrix (denoted as *M*) with rows corresponding infused metabolites and columns to measured metabolites. The entry (*i, j*) of *M* is the normalized labeling of metabolite *j* by metabolite *i*. The diagonal entries of the matrix represent the normalized labeling of the infused metabolite itself and are 1 by the definition of normalized labeling. Experimental data for the 15 circulating metabolites studied here (the 13 highest flux individual metabolites + the 2 essential amino acid mixtures) are shown in Table S2.

The direct contribution to circulating metabolite *k* from the other 14 circulating metabolites is given by a system of 14 linear equations:

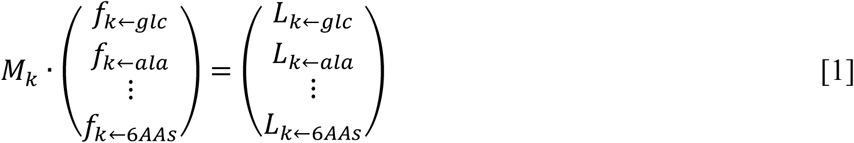

*M*_*k*_ is the 14 × 14 submatrix of *M* with both the row and column corresponding to *k* removed, *L* is the column vector of the normalized labeling data from the isotope infusion experiments (thus the *k-th* column of *M* with the *k-th* entry removed), and *f* is the length 14 column vector of direct contributions. While the equations can be solved for the direct contributions, to avoid negative values due to measurement errors, we use an optimization to obtain the direct contributions (see Methods). Application of this procedure to lactate revealed that infusions of glucose, alanine, and other amino acids label lactate, but glucose is the predominant direct contributor: the amino acids are first converted to circulating glucose by gluconeogenesis and then the resulting labeled glucose makes lactate (Figure 2B). This discrepancy emphasizes the importance of determining direct contributions for elucidating the *in vivo* operation of metabolism.

### Circulating Metabolite Sources

Eqn. 1 can be applied to each of the 15 nutrients to determine its direct contributions from the other 14 nutrients. The comprehensive direct contribution analysis in 8-h fasted mice reveals fundamental aspects of metabolism (Figure 2C; Table S3). Circulating glucose, lactate, alanine and 3-hydroxybutyrate come mainly from other circulating metabolites, as the sum of the circulating metabolite contributions is almost 1. In contrast, the circulating metabolite contributions to fatty acids and essential amino acids are negligible, reflecting their production instead by hydrolysis of triglycerides and protein. Circulating glucose is made from a roughly equal balance of lactate, alanine, and glycerol. Glucose is the dominant precursor of lactate, and lactate the main source of alanine. Circulating glycerol is made from both glucose and lactate, but mainly comes from stored triglycerides. Acetate is produced roughly equally from circulating metabolites and other sources (e.g. the microbiome, protein deacetylation). Despite recent evidence that glucose catabolism can make acetate (Liu et al., 2018), the predominant circulatory precursors to acetate are fatty acids.

### Absolute Fluxes between Circulating Metabolites

The direct contribution analysis reveals the existence of futile metabolic cycles. For example, consistent with the Cori cycle, there is direct contribution from circulating lactate to circulating glucose, and vice versa (Figure 2C). While cycles can be identified from the direct contributions, to quantify the magnitude of the cycles, it is necessary to know the absolute fluxes between circulating metabolites. We denote the flux from circulating metabolite *i* to metabolite *k* as *J*_*k*←*i*_. Despite decades of research into metabolic cycles, a general method for determining such absolute fluxes has been lacking. Intuitively, the flux from *i* to *k* should reflect how fast circulating *k* is being produced (conceptually, *F*_*circ_k*_) multiplied by the fraction of *k* coming from *i* (*f*_*k*←*i*_). The challenge is that *F*_*circ*_ (and equivalently classical measurements of *R*_*a*_) actually measure dilution of the infused isotope-labeled metabolite in the bloodsteam by incoming flux from diet or synthesis. If the incoming flux from synthesis is in part labeled (due to metabolic cycling), the absolute magnitude of the flux will be underestimated.

We developed an algorithm that uses the comprehensive interconversion data (i.e., the direct contributions between metabolites in Figure 2C and Table S3) to calculate a conversion factor (denoted as *c*_*k*_) between the isotope measured *F*′_*circ,k*_ and the absolute circulatory flux through that metabolite (*J*_*circ,k*_). Absolute fluxes connecting circulating metabolites can then be determined by multiplying *J*_*circ,k*_ by the direct contribution values (Figure 2D). Details of the algorithm can be found in Suppl. Note. The biggest absolute fluxes are glucose to lactate and lactate, alanine, and glycerol to glucose (Figure 2E).

### Cycling of Circulating Nutrients

We define the magnitude of a metabolic cycle among nutrients as the minimum of the inter-converting fluxes between the nutrients, 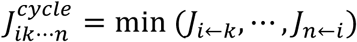. Figure 2F shows all metabolic cycles between circulating metabolites with flux greater than 50 nmol carbon/min/g (about 10% of the glucose *F*_*circ*_). This analysis identified three highest flux metabolic cycles: glucose-lactate-glucose, lactate-alanine-lactate, and glucose-lactate-alanine-glucose (Figure 2F). This latter pathway is particularly interesting, reflecting that lactate (not glucose) is the predominant source of circulating alanine, and that alanine is co-equal with lactate as a contributor to gluconeogenesis, despite having much lower *F*_*circ*_. The analysis also identified a previously unrecognized metabolic cycle between circulating lactate and glutamine, reflecting the importance of lactate as a TCA substrate and substantial efflux from the TCA cycle to lactate.

### Persistence of the Metabolic Cycling in Fed Mice

We have so far focused on carbohydrate-diet mice fasted for 8 h (9 AM to 5 PM, when they sleep and food intake is typically low). Over this duration, the respiratory exchange ratio (RER) drops from above 0.9 (reflecting mainly carbohydrate burning) to less than 0.8 (reflecting mainly fat burning) (Figure 3A). As expected, refeeding increases the RER consistent with resumption of carbohydrate burning (Figure 3A). Despite the strong change in RER, comprehensive *F*_*circ*_ measurement revealed strikingly similar fluxes between fasted and refed mice, with the exception of much higher *F*_*circ_glucose*_ in the fed state, reflecting glucose influx from the meal (Figure 3B; Table S4). The persistent lactate and fatty acid fluxes suggest continued Cori cycling and triglyceride cycling in the fed state.

**Figure 3.**
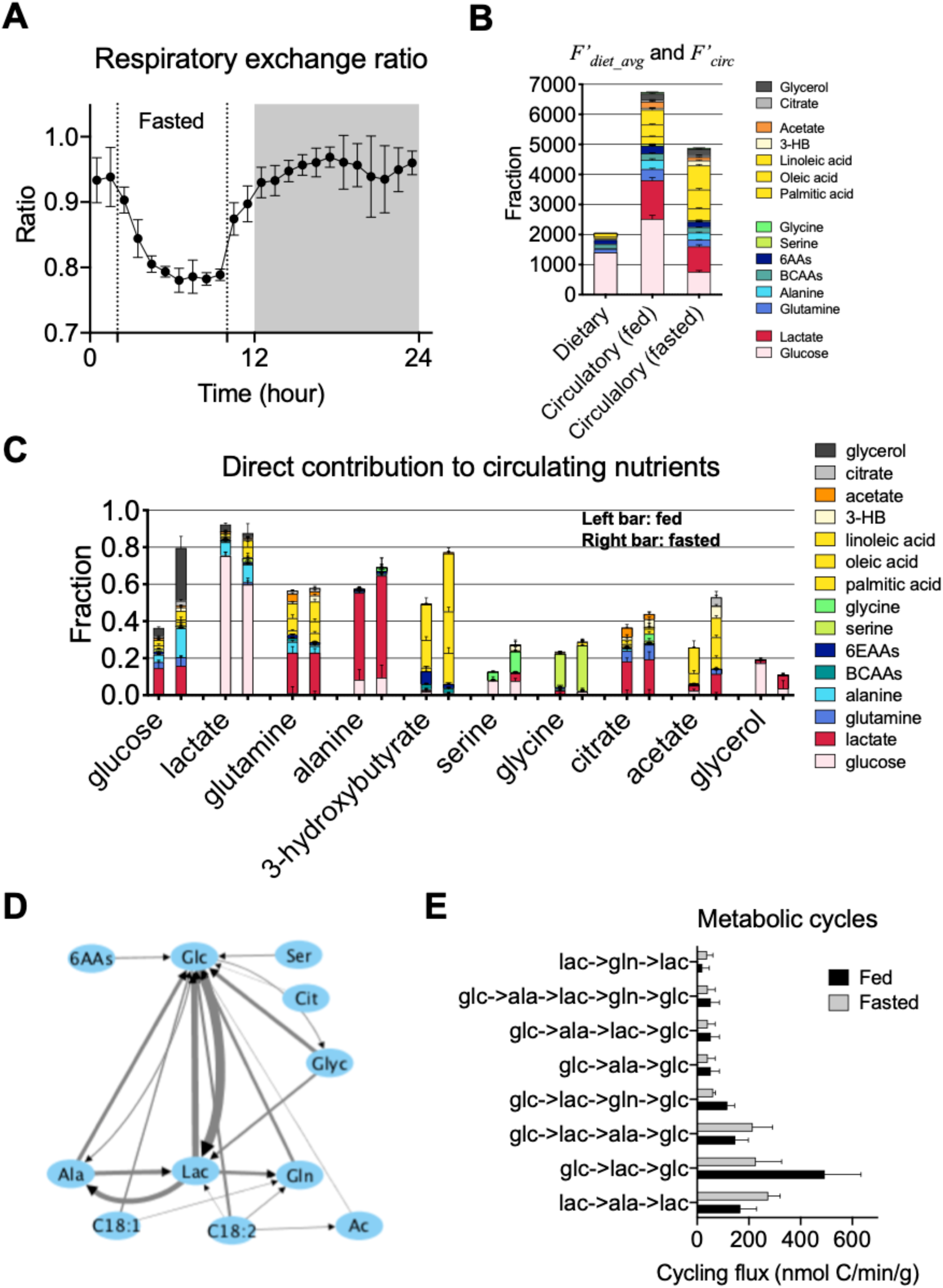
Metabolite interconversion fluxes are similar between fed and fasted mice. Data are for 8-h fasted mice and 3-h refed mice on carbohydrate diet. (A) Respiratory exchange ratio during a diurnal cycle (N=5). Shaded area indicates the dark period. Mice were 8-h fasted during the light period. (B) Dietary flux versus circulatory turnover flux in fed and fasted mice. See Tables S1 and S4 for flux values and N. (C) Direct contributions to circulating metabolites in both fed and fasted mice. For each metabolite, the left bar is fed state and right bar is fasted. See Table S2 for normalized labeling and Tables S3 and S5 for direct contribution values. (D) Circulating nutrient interconversion fluxes in fed mice. Edge widths are proportional to log-transformed flux values. Shown are fluxes > 50 nmol C/min/g. (E) Top metabolic cycles in fed and fasted mice.

We then proceeded to analyze direct contributions to circulating metabolites in the fed state. For many metabolites, including lactate, alanine, and glutamine, their sources were nearly unaltered. For others, including glucose, serine, and acetate, the fraction produced from circulating metabolites fell with feeding, reflecting their coming instead from the diet, or, in the case of acetate, the gut microbiome (Figure 3C; Tables S3 and S5). Analysis of absolute fluxes connecting circulating metabolites revealed no substantial new fluxes in the fed state. Metabolic cycling centered around glucose, lactate, and alanine was, however, persistent (Figure 3D,E).

### Measuring Direct TCA Cycle Contributions

A primary function of circulating metabolites is to support tissue energy production, which occurs primarily through the TCA cycle. We next aimed to determine the direct contribution of any circulating metabolite *X* to the TCA cycle of different tissues. The experimental approach is to infuse uniformly ^13^C-labeled *X* and measure succinate and malate labeling. These two TCA intermediates are chosen for their higher concentration in tissues than in serum (Figure S1), enabling robust tissue-specific measurements. We define the normalized TCA labeling of circulating metabolite *X* to the TCA cycle of tissue 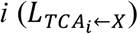 as the fraction of labeled carbon atoms in tissue malate or succinate relative to that of *X* in serum.

Due to the interconversion of circulating nutrients, the normalized TCA labeling of *X* may exceed its direct contribution, with *X* instead feeding the TCA cycle via other circulating metabolites (Figure 4A). For example, both glucose and glycerol infusions label TCA intermediates in most tissues (Figure 4B), but such labeling may occur indirectly via circulating lactate (Hui et al., 2017). The measured TCA labeling reflects the weighted sum of the direct and indirect routes from the infused tracer to the tissue TCA. For example, in Figure 4A, the measured TCA labeling by *X* 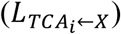 can be expressed in terms of the normalized labeling of *Y* (*L*_*Y*←*X*_) and *Z* (*L*_*Z*←*X*_) and the three direct contributions to TCA from *X, Y*, and *Z* (denoted as 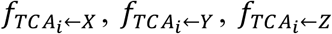, respectively). To determine all the three direct contributions in this example, we also need to infuse the other two nutrients and measure the labeling of both tissue malate and the other two nutrients in serum. The resulting three equations can then be used to solve for the three unknowns.

**Figure 4.**
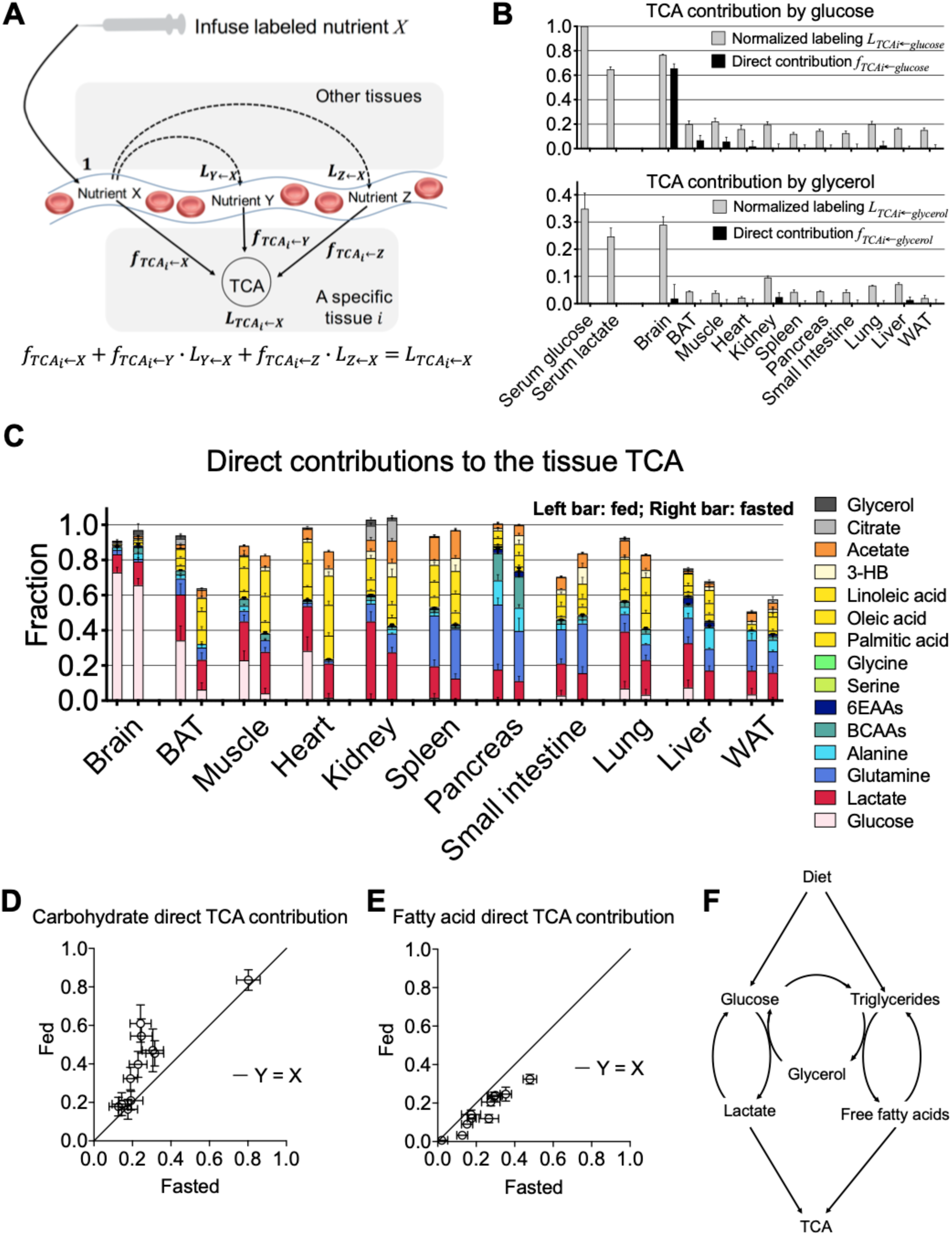
Comprehensive tracer studies enable determination of direct sources of tissue TCA metabolites. Data are for 8-h fasted mice and 3-h refed mice on carbohydrate diet. (A) Example of direct and indirect routes from the infused tracer to tissue TCA cycle. The labeling of the tissue TCA upon labeled *X* infusion can come from both directly from *X* or indirectly via circulating *Y* or *Z*, as codified in the isotope-balance equation. Infusions of *Y* and *Z* yield analogous equations which are needed to solve for the three direct contributions. (B) Comparison between the normalized labeling of tissue TCA intermediate (malate) and direct contribution for glucose (upper panel) and glycerol (lower panel). (C) Direct tissue TCA contributions for 15 circulating nutrients and 11 tissues. For each tissue, the left bar is fed state and right bar is fasted. See Table S2 for normalized labeling and Table S6 for direct TCA contribution values. (D) Comparison of direct contribution from carbohydrates (the sum of direct contributions from glucose and lactate) to tissue TCA between fasted and fed mice. (E) Comparison of direct contribution from fatty acids (the sum of direct contributions from palmitic acid, oleic acid, and linoleic acid) to tissue TCA between fasted and fed mice. (F) Schematic of major carbohydrate and fatty acid fluxes: constant functioning of the glucose-lactate cycle and the triglyceride-glycerol-fatty acid cycle, with glycerol connecting the two, and lactate and free fatty acids the primary direct tissue TCA contributors.

We generalized this scheme to determine the direct TCA contributions of the 13 high flux circulating metabolites plus the two amino acid mixtures. To this end, in addition to measuring the interconversion matrix *M*, we also measured malate and succinate labeling from each of the 15 tracers across 11 tissues. For each tissue, the direct TCA contributions 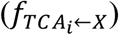 are determined by solving 15 linear equations:

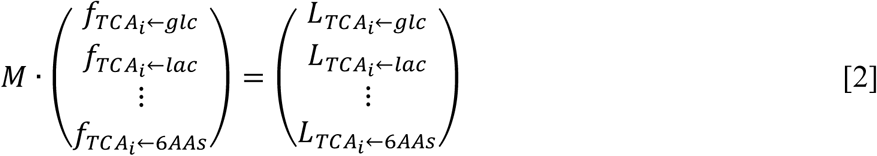

As in the case of calculating direct contributions between circulating nutrients, to avoid negative values due to measurement errors, we use an optimization to obtain the direct contributions (see Methods). In the case of glucose and glycerol, direct TCA contributions are much lower than measured TCA labeling, emphasizing the capacity for this quantitative analysis to unveil metabolic fundamentals that are normally masked by interconversion among circulating nutrients (Figure 4B).

### TCA Fuel Choices across Tissues

The tissue-specific direct TCA contributions provide a systems-level perspective of fuel utilization in both 8-h fasted and refed mice (Figure 4C; Table S6). The same trends are obtained using either malate or succinate labeling as the readout, supporting the robustness of the results (Figure S2). Brain is unique in mainly using glucose in both the fed and fasted states. In contrast to literature suggesting that the heart is an omnivore (Bing et al., 1954; Drake et al., 2012), cardiac muscle stands out for barely using amino acids as fuels, preferentially consuming fatty acids. Kidney is unique in burning circulating citrate (Jang et al., 2019). Pancreas shows the greatest use of amino acids.

Consistent with the comprehensiveness of our analysis, in most tissues, the measured direct contributions almost add up to 1, reflecting a nearly comprehensive accounting for TCA carbon inputs. Notable exceptions are adipose and liver, where the missing carbon likely comes from burning of internal nutrient stores. In the small intestine, the missing contribution may also include direct input from diet.

In general, tissues differed markedly in their metabolic preferences, and maintained these preferences across the fed and fasted states. This manifests as a strong correlation between fasted and fed tissue carbohydrate (glucose/lactate) and fat usage (Figure 4D, E). In addition, there is a shift towards a greater carbohydrate contribution in feeding, and fat contribution in fasting (Figure 4D, E), with brown adipose and muscle being particularly responsive (Figure 4C).

The analysis also reveals which circulating metabolites directly drive TCA metabolism, with lactate, fatty acids, and (to a lesser extent) acetate serving as major contributors in nearly all tissues (Figure 4B). Glucose is the dominant precursor of circulating lactate (Figure 3C), but itself contributes directly to TCA in only selected instances. Similarly, glycerol carries substantial circulatory flux and its infusion labels TCA in most tissues (Figure 4B), but it makes essentially no direct contribution (Figure 4C). Instead of being burned in TCA, glycerol is a dedicated gluconeogenic precursor (Figure 2C).

Together, the *F*_*circ*_, metabolite interconversion, and TCA contribution data support a model in which circulating glucose is constantly being interconverted with circulating lactate, and free fatty acids are constantly being released from and stored in triglycerides, with the bulk of energy being generated by tissue uptake and burning of lactate and free fatty acids, rather than glucose or triglycerides themselves (Figure 4F).

### Persistence of the Metabolic Cycling in Mice on Ketogenic Diet

Mammals cannot synthesize certain amino acids and fatty acids, and hence these are essential dietary components. Given adequate supply of these essential nutrients, mammals can survive on nearly pure carbohydrate or pure fat as the dominant caloric input. Standard lab chow, like a modern human diet high in bread or rice, provides most calories as carbohydrate. An opposite extreme is ketogenic diet, which is mainly fat with minimal carbohydrate (25-fold lower than standard lab chow; Table S7). Ketogenic diet has been successfully used for almost a century as a treatment for refractory epilepsy (Barañano and Hartman, 2008; D’Andrea-Meira et al., 2019; Lima et al., 2014). More recently, ketogenic diet is attracting increasing interest for its ability to ameliorate type II diabetes (McKenzie et al., 2017), and even slow the progression of certain cancers (Allen et al., 2014; Hopkins et al., 2018; Lussier et al., 2016; Mavropoulos et al., 2009; Otto et al., 2008).

To examine how a radical change of diet affects metabolism, we performed both classical metabolic phenotyping (Figure S3) and circulatory flux analyses on mice on a ketogenic diet. Consistent with literature, ketogenic diet slightly decreased body weight and increased whole body oxygen consumption and energy expenditure (Jornayvaz et al., 2010; Kennedy et al., 2007; Pissios et al., 2013). Ketosis is confirmed with the elevated serum 3-hydroxybutate level on the ketogenic diet (Figure S4). Despite the dietary inputs being almost completely different (Figure 5A; Table S8), *F*′_*circ*_ values were similar to standard carbohydrate diet (Figure 5B; Table S8). This supports the model in which chronic glucose-lactate cycling and triglyceride cycling renders internal metabolic activity robust to changing dietary inputs.

**Figure 5.**
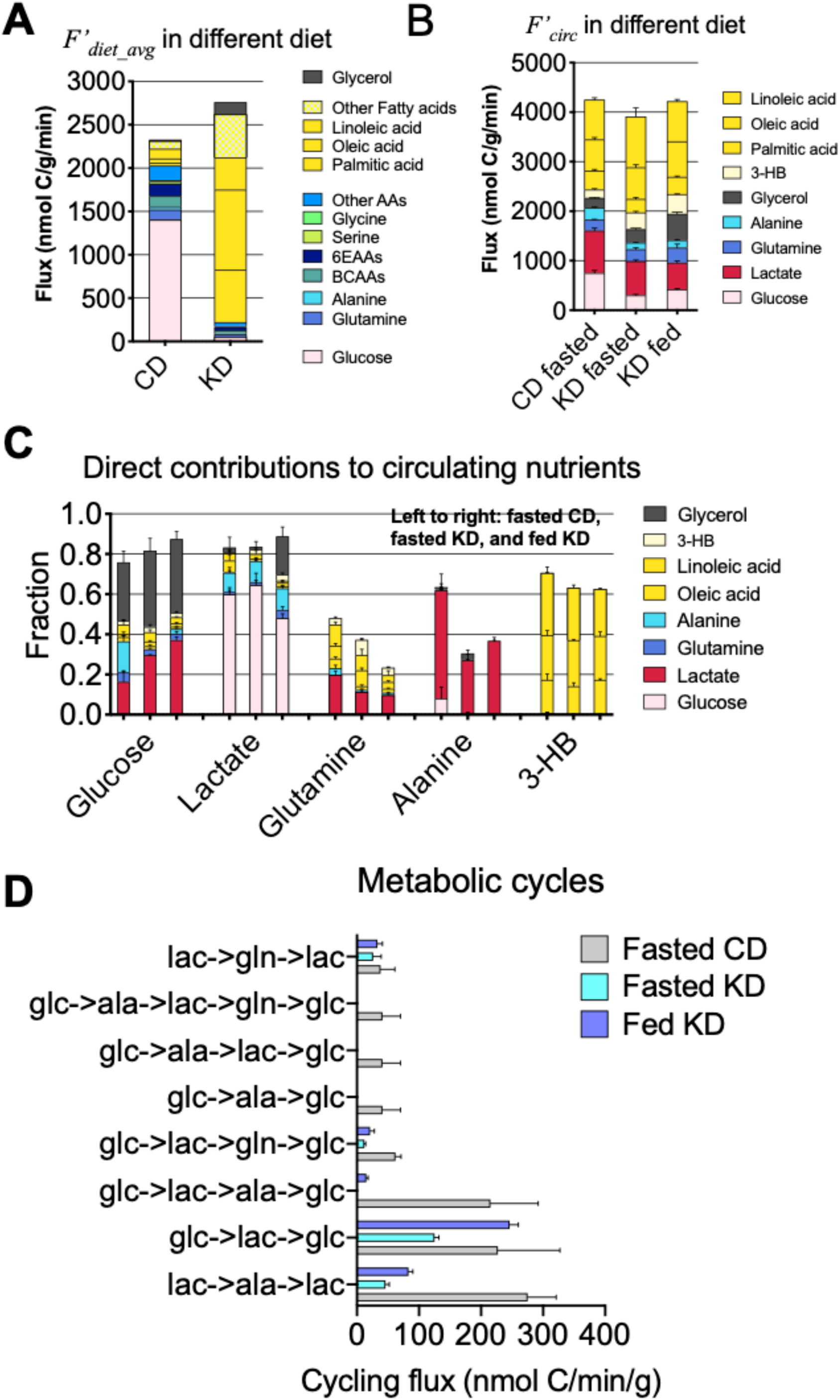
Persistent circulatory carbohydrate fluxes on ketogenic diet. Data are for 8-h fasted carbohydrate diet mice and 8-h fasted and 3-h refed ketogenic diet mice. (A) Comparison of the dietary fluxes. See Tables S1 and S7 for flux values and N. (B) Comparison of the circulatory turnover fluxes. See Table S1 and S7 for flux values and N. (C) Comparison of the direct contributions to circulating metabolites. For each metabolite, the left bar is fasted carbohydrate diet, middle bar is fasted ketogenic diet, and right bar is fed ketogenic diet. See Table S8 for normalized labeling and Table S9 for direct contribution values. (D) Comparison of metabolic cycles between the two diets.

To explore this concept further, we infused nine tracers (Table S9) to determine their interconversions and TCA inputs in ketogenic diet (Figure 5C; Table S10). Alanine disappeared as a major gluconeogenic precursor in the ketogenic diet, while glycerol emerged as a direct lactate precursor. Despite these shifts, the primary sources of each circulating metabolite remained the same. Moreover, while metabolic cycling involving alanine decreased, glucose-lactate cycling persisted, despite the near complete lack of dietary carbohydrate (Figure 5D). Thus, the nutrient cycling persists regardless of the dietary intake.

### Pyruvate Cycling in Ketogenic Diet

Consistent with the low dietary intake of carbohydrates, the RER for mice on the ketogenic diet remained low (∼0.73) throughout the diurnal cycle (Figure 6A), indicating predominantly fatty acid burning. Assessment of TCA substrates revealed extensive fat and much decreased carbohydrate utilization in skeletal and cardiac muscle and brown adipose tissue (Figure 6B; Table S11). In addition, 3-hydroxybutyrate, which is derived from fatty acids (Figure 5C), emerged as a broadly important direct TCA input (Figure 6B). Interestingly, despite the paradigm that brain shifts from using glucose to primarily using 3-hydroxybutyrate during ketosis (LaManna et al., 2009; Zhang et al., 2013), glucose remained the largest brain TCA substrate, with lactate serving as also a major contributor. Collectively, even on ketogenic diet, more than 60% of brain TCA carbon still comes from carbohydrate (Figure 6C). Similarly, there was a substantial persistent lactate contribution to TCA cycle in many tissues, most notably kidney and liver (Figure 6C). Thus, analysis of TCA inputs revealed persistent carbohydrate metabolism in brain, kidney, and liver.

**Figure 6.**
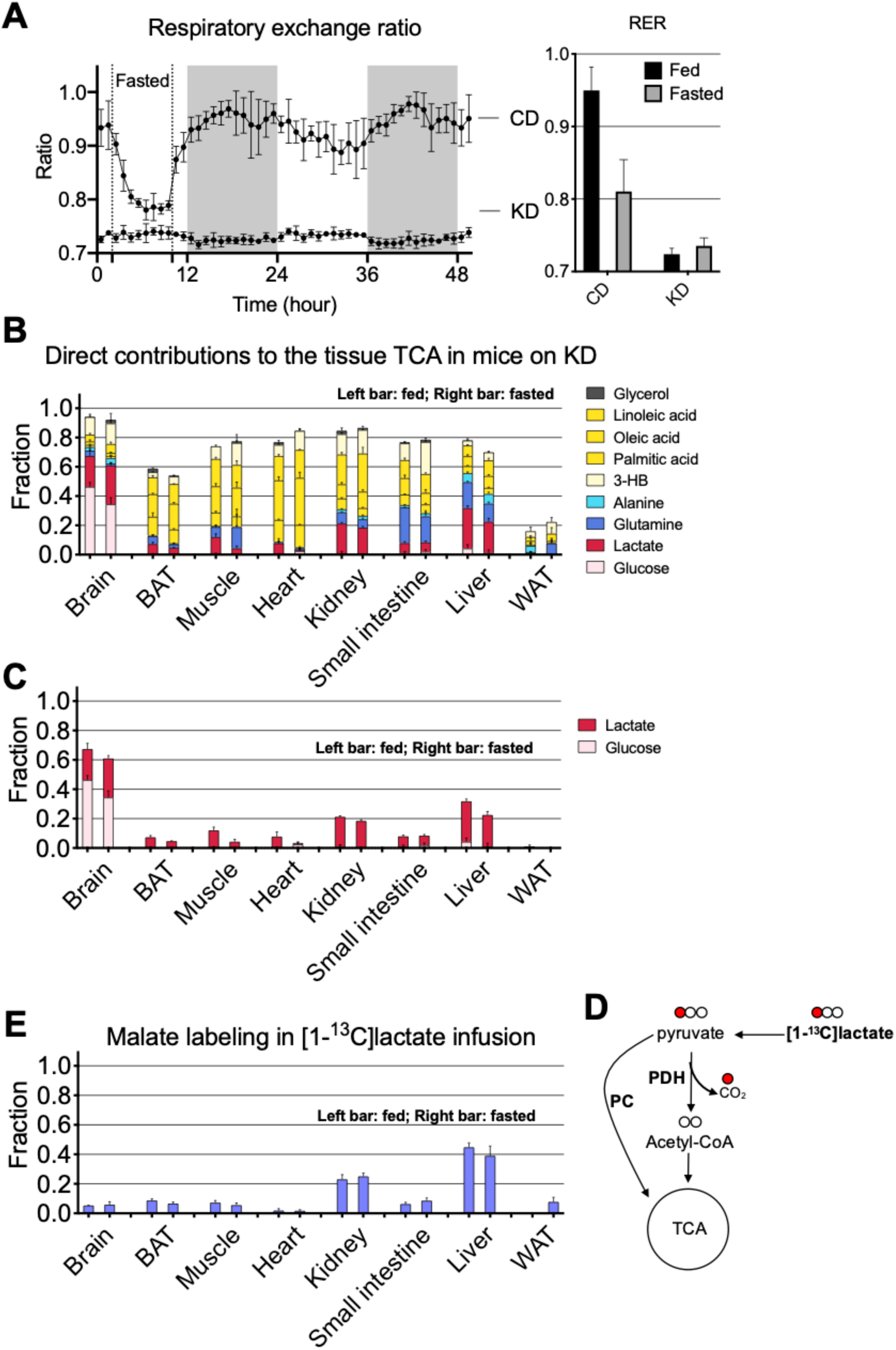
In ketogenic diet, pyruvate cycling produces carbohydrate fluxes without carbohydrate burning. (A) Comparison of respiratory exchange ratio between mice on the high-carbohydrate diet (N=5) and the ketogenic diet (N=4). Shaded areas indicate the dark periods. Mice were fasted for 8-hr in the first light period. Error bars are standard deviations. (B) Direct contributions from 9 circulating metabolites to the TCA cycle in tissues of fasted mice and fed mice on the ketogenic diet. See Table S8 for normalized labeling and Table S10 for direct TCA contribution values. (C) Same data as in (B), but only the glucose and lactate data are shown. (D) The labeled carbon of [1-^13^C]lactate is lost in the PDH reaction, and hence labeling observed in (D) reflects TCA input via the PC pathway. (E) Analogous to (C), with [1-^13^C]lactate as the tracer.

Carbohydrate can enter the TCA cycle through two routes: oxidation to acetyl-CoA by pyruvate dehydrogenase (PDH) or carboxylation to oxaloacetate. The former burns carbohydrate, while the latter generates four-carbon units that can remake glucose via gluconeogenesis. PDH removes C1 of lactate/pyruvate (which comes from C3/C4 of glucose). Hence, [1-^13^C]lactate infusion can be used to selectively trace TCA entry via pyruvate carboxylase (Figure 6E). Strikingly, on ketogenic diet, the persistent carbohydrate TCA contribution in kidney and liver reflects strong pyruvate carboxylase activity. Such activity, which is a general feature of many tissues (Figure 6D), does not burn carbohydrate, but rather supports glucose-lactate cycling. Extensive pyruvate carboxylase activity in kidney and other non-hepatic organs implies that analysis of pyruvate cycling needs to consider the whole body, not solely liver metabolism (Burgess et al., 2015; Hasenour et al., 2015; Perry et al., 2016). Notably, the large carbohydrate contribution to brain TCA on ketogenic diet is not associated with substantial pyruvate carboxylase activity, and instead reflects carbohydrate burning. Collectively, these observations support a model in which glucose-lactate cycling persists in ketogenic diet, driven by lactate uptake into both liver and kidney TCA via pyruvate carboxylase to support gluconeogenesis, with carbohydrate burning via pyruvate dehydrogenase turned off in all organs except for the brain.

## DISCUSSION

Mammals are nourished by the food we eat. Food is initially converted into monomeric nutrients like glucose and amino acids. These nutrients can then be stored or interconverted. This processing serves to ensure that the circulation carries a steady supply of appropriate nutrients to each tissue, despite varying dietary intake. To examine this processing comprehensively and quantitatively, we carried out minimally perturbative isotope-tracer infusions for all high flux circulating metabolites. Using LC-MS, we measured experimentally the extent to which any given metabolite labels other circulating metabolites, as well as its tissue-specific TCA contributions.

The comprehensiveness of these experiments enables quantitative analyses that are not possible from studying any single nutrient in isolation. Specifically, the experimentally observed labeling from any given nutrient tracer reflects the sum of all routes linking the tracer to the measured metabolic product. Accordingly, the direct metabolic connections are clouded by interconversions among nutrients. Using our comprehensive measurements, through robust and readily implemented linear algebra analyses, we illuminate the direct metabolic connections and their quantitative magnitudes. In doing so, we provide fundamental reference data regarding *in vivo* metabolic activity and methods that can be applied across organisms, diets, and disease states.

Glycerol metabolism highlights the utility of determining direct contributions, rather than merely labeling. We find that glycerol is unique among high-flux circulating nutrients in not being catabolized into the TCA cycle in any tissue. This makes physiological sense. To survive prolonged starvation, mammals catabolize stored triglycerides, which yields fatty acids and glycerol. Because fatty acids cannot support gluconeogenesis, the resulting three carbon units are precious and there is no sense in burning glycerol. Importantly, however, a naïve labeling measurement would show a clear TCA contribution from glycerol, due to its indirect TCA contributions via circulating glucose and lactate. Its special physiological role becomes clear from our systems-level analysis.

Direct contributions also provide a sharp picture of tissue TCA fuel preferences. Muscle and brown adipose show strong responses to feeding, directing glucose carbon directly into their TCA cycle selectively in the fed state. In contrast, most other tissues show remarkably stable fuel preferences across feeding, fasting, and even ketogenic diet. Lactate is unique in being a universal TCA substrate. Fat is also widely used, except in brain. Acetate is a relatively major contributor in many organs, but not brain or liver where it is instead directed towards lipogenesis (Zhao et al., 2016). Amino acids are preferred fuels for the visceral organs but not muscle.

Nutrient interconversion enables tissues to access their preferred fuels, even when dietary content changes. In the United States, roughly half of caloric intake is carbohydrates, one-third fat, and one-sixth protein (CDC, 2000). Mammals can thrive, however, eating either minimal fat or minimal carbohydrate (Hu et al., 2018; Simpson and Raubenheimer, 2012). Diets sufficiently low in both carbohydrate and protein are ketogenic: they lead to high circulating levels of ketone bodies, most importantly 3-hydroxybutyrate (3HB), which are made from fat by liver. Given that the brain does not effectively take up fatty acids, 3HB is widely assumed to be the major brain fuel during ketosis (LaManna et al., 2009; Zhang et al., 2013). Our analyses paint a different picture. 3HB contributes broadly to TCA, to a similar extent in brain, muscle, kidney, and intestine. In the brain, glucose and lactate, the two main forms of circulating carbohydrate, remain greater TCA contributors than 3HB. The continuous supply of carbohydrate to the brain, despite its near absence in the diet, is enabled by conversion of the glycerol moiety in dietary triglycerides into circulating glucose by the liver and kidney.

A key finding of our work is that such metabolic interconversions do not occur at a minimal rate, just sufficient for tissue demands. Instead, we observe rapid futile cycling, in particular of nutrients that are scarce in the diet. For example, in standard high carbohydrate chow, we observe high circulatory turnover fluxes of fatty acids, with circulatory fatty acid and carbohydrate fluxes of similar magnitude. This reflects continuous triglyceride breakdown, release of fatty acids into the bloodstream, and their subsequent uptake and reincorporation into tissue fat. This cycling ensures a robust supply of fat and glycerol should demand arise (Newsholme et al., 1983). Strikingly, in ketogenic diet, circulatory fatty acid fluxes are no higher than in the high carbohydrate diet, and circulatory carbohydrate fluxes are only modestly lower. This reflects continuous synthesis of lactate from glucose and glucose from lactate. Alanine fluxes are, however, down, presumably due to lower dietary protein intake decreasing the need for nitrogen shuttling to the liver.

The relatively invariant circulatory fluxes of fatty acids, glycerol, lactate, and glucose across diets does not mean that the importance of these nutrients as fuels is steady. To maintain homeostasis, mammals must burn what they eat. This divergence between nutrient cycling, which is relatively insensitive to diet, and nutrient burning, which changes with diet, is neatly captured by tracing positionally labeled lactate into the TCA cycle. In ketogenic diet, lactate continues to enter the TCA cycle, especially in brain, liver and kidney. In brain, this entry is via PDH, reflecting brain continuing to burn carbohydrate. In liver and kidney, however, entry is via pyruvate carboxylase, which does not destroy the three-carbon unit, but instead initiates its use for gluconeogenesis. Functionally, this makes sense. Persistent nutrient cycling ensures availability of the most important energy substrates to tissues irrespective of the physiological state. Tight regulation of burning ensures that limiting nutrients are saved for the tissues that need them most.

Overall, we lay out quantitative, systems levels methods for dissecting mammalian metabolism. The heart of these methods is mathematical integration of diverse tracer data to reveal direct connections both among circulating metabolites and between circulating metabolites and tissue TCA intermediates. Direct connections are not the only ones that matter functionally. For example, most lactate carbon originates from glucose. Accordingly, the minimal direct TCA contribution of glucose does not reflect a lack of glucose burning, but merely that circulating lactate is an intermediate in this process. In this way, direct contributions provide discrete measures of metabolic activity that can be used to advance quantitative understanding of metabolic physiology. Going forward, it will be important to relate these quantitative physiological measures both to underlying biochemical events and to the action of hormonal regulators that control metabolic health.

## STAR*METHODS

### LEAD CONTACT AND MATERIALS AVAILABILITY

Further information and requests for resources and reagents should be directed to and will be fulfilled by the Lead Contact, Joshua D. Rabinowitz.

### EXPERIMENTAL MODEL AND SUBJECT DETAILS

Mouse studies followed protocols approved by the Animal Care and Use Committee for Princeton University and for the University of Pennsylvania. *In vivo* infusions were performed on 12-14 week old C57BL/6 mice pre-catheterized in the right jugular vein (Charles River Laboratories, Wilmington, MA). Animals received either a normal chow diet (PicoLab Rodent 20 5053 laboratory Diet St. Louis, MO) or a ketogenic diet (Bio-Serv F3666 Flemington, NJ). Mice were allowed at least 5 days of acclimation to the facilities prior to experimentation and were randomly chosen for infusions of different tracers. No blinding was implemented. Those animals receiving the ketogenic diet were adapted to the diet for at least 21 days before experimentation. The mice were on normal light cycle (8 AM – 8 PM). On the day of infusion experiment, mice were transferred to new cages without food around 9 AM (beginning of their sleep cycle) and infused for 2.5 h starting at around 3 PM. To probe the fed state, the mice were maintained without food until around 8 PM, at which time chow was placed back in the cages and the 2.5 h infusion initiated. The infusion setup (Instech Laboratories) included a swivel and tether to allow the mouse to move around the cage freely. Water-soluble isotope-labeled metabolites (Cambridge Isotope Laboratories, Tewksbury, MA) were prepared as solutions in sterile normal saline. To make ^13^C-labeled fatty acid solutions, the fatty acids were complexed with bovine serum albumin in a molar ratio 4:1. Infusion rate was set to 0.1 μl min^-1^ g^-1^ for water-soluble metabolites and 0.4 µl min^-1^ g^-1^ for fatty acids. Blood was collected by tail snip (∼10 μl) and transferred into blood collection tubes with clotting factor (Sarstedt 16.442.100). Blood samples were stored on ice and then centrifuged at 16,000 x g for 10 minutes at 4°C to get serum samples. Tissue harvest was performed at the end of the infusion after euthanasia by cervical dislocation. Tissues were quickly dissected, clamped with a pre-cooled Wollenberger clamp, and dropped in liquid nitrogen.

## METHOD DETAILS

### Metabolite Extraction of Serum

Serum (2 μl) was added to 68 μl of -80°C 100% methanol, vortexed, and put on dry ice for at least 5 minutes. Following, extract was centrifuged at 16,000 x g for 10 minutes at 4°C and supernatant was mixed 1:1 with 80% methanol. After centrifugation again at 16,000 x g for 10 minutes at 4°C, supernatant was transferred to tubes for LC-MS analysis.

### Metabolite Extraction of Tissue

Frozen tissue was ground by a Cyromill at cryogenic temperature (Retsch, Newtown, PA). Ground tissue was then weighed (∼20mg) and mixed with -20°C 40:40:20 methanol:acetonitrile:water with 0.5% formic acid (extraction solvent) at a concentration of 25mg/ml. Samples were briefly vortexed before neutralizing with 8 μl of 15% ammonium bicarbonate per 100 μl of extraction solvent. Extract was then vortexed and centrifuged twice at 16,000 x g for 20 minutes at 4°C before the final supernatant was transferred to LC-MS tubes for analysis.

### Metabolite Measurement by LC-MS

A quadrupole-orbitrap mass spectrometer (Q Exactive, Thermo Fisher Scientific, San Jose, CA) operating in negative mode was coupled to hydrophilic interaction chromatography (HILIC) via electrospray ionization. Scans were performed from m/z 70 to 1000 at 1 Hz and 140,000 resolution. LC separation was on a XBridge BEH Amide column (2.1 mm x 150 mm x 2.5 μm particle size, 130 Å pore size; Water, Milford, MA) using a gradient of solvent A (20 mM ammonium acetate, 20 mM ammounium hydroxide in 95:5 water:acetonitrile, pH 9.45) and solvent B (acetonitrile). Flow rate was 150 μl/min. The LC gradient was: 0 min, 85% B; 2 min, 85% B; 3 min, 80% B; 5 min, 80% B; 6 min, 75% B; 7 min, 75% B; 8 min, 70% B; 9 min, 70% B; 10 min, 50% B; 12 min, 50% B; 13 min, 25% B; 16 min, 25% B; 18 min, 0% B; 23 min, 0% B; 24 min, 85% B. Autosampler temperature was 5°C, and injection volume was 5-10 μl for serum samples and 15 μl for tissue samples. For improved detection of fructose-1,6-bisphosphate and 3-phosphoglycerate, selected ion monitoring (SIM) scans were added. Data were analyzed using the El-MAVEN software. For tracer experiments, isotope labeling was corrected for natural ^13^C abundance.

Circulating glycerol labeling was determined by first converting serum glycerol to glycerol-3-phosphate using glycerol kinase and then measuring the labeling of glycerol-3-phosphate with LC-MS.

Acetate was derivatized and measured by LC-MS. The derivatizing reagent was 12 mM EDC, 15 mM 3-Nitrophenylhydrazine and pyridine (2% v/v) in methanol. Reaction was stopped with quenching reagent consisting of 0.5 mM beta-mercaptoethanol and 0.1% formic acid in water. Serum (5 µL) was mixed with derivatizing reagent (100 µL) and incubated for 1 hour at 4°C. Then, the samples were centrifuged at 16,000 x g for 10 min at 4°C, and 20 µL of supernatant was mixed with 200 µL of the quenching reagent. After centrifugation at 16,000 x g for 10 min at 4°C, supernatants were collected for LC-MS analysis. A quadrupole-time of flight mass spectrometer (Q-TOF, Agilent, Santa Clara, CA) operating in negative ion mode was coupled to C18 chromatography via electrospray ionization and used to scan from m/z 100 to 300 at 1 Hz and 15,000 resolution. LC separation was on an Acquity UPLC BEH C18 column (2.1 mm x 100 mm, 1.7 5 µm particle size, 130 Å pore size; Waters, Milford, MA) using a gradient of solvent A (water) and solvent B (methanol). Flow rate was 200 µL/min. The LC gradient was: 0 min, 10% B; 1 min, 10% B; 5 min, 30% B; 7 min, 100% B; 11 min, 100% B; 11.5 min, 10% B; 14 min, 10% B. Autosampler temperature was 5°C, column temperature was 60°C and injection volume was 10 µL. Ion masses for derivatized acetate was 194.

### Body Composition and Indirect Calorimetry

Body weight and body composition were measured in ad-lib fed conscious mice using EchoMRI™ 3-in-1 system nuclear magnetic resonance spectrometer (Echo Medical Systems, Houston, TX) to determine whole body lean and fat mass. Indirect calorimetry was performed in individually housed mice using a 20-channel open-circuit indirect calorimeter, in which 10 cages were mounted inside two thermally controlled cabinets, maintained at 22°C (Comprehensive Lab Animal Monitoring System; Columbus Instruments, Columbus, OH, USA). After overnight acclimation (20 hours), oxygen consumption (VO2), the amount of carbon dioxide produced (VCO2), energy expenditure, food and water intake, ambulatory and locomotor activity (infrared beam breaks) were determined during an initial period of fasting (9am-5pm), as well as for the following 40 hours wherein mice were fed *ad libitum*. Average respiratory exchange ratio (RER) was calculated as the ratio of VCO2:VO2 over 48 hours. Energy expenditure was calculated as heat (kcal/hr)=(3.815 +1.232 x RER) x VO2). The metabolic studies were performed at the Penn Diabetes Research Center Rodent Metabolic Phenotyping Core (University of Pennsylvania).

## QUANTIFICATION AND STATISTICAL ANALYSIS

### Calculation of average dietary fluxes

The indirect calorimetry determined that the energy expenditure was 0.39±0.02 Calories/day/g for mice on the carbohydrate diet (CD) and 0.53±0.04 Calories/day/g for mice on the ketogenic diet (KD). Using the calorie density for the diets (3.41 Calories/g for CD and 7.24 Calories/g for KD), we have the total dietary fluxes as 0.115 g CD/g of mouse/day and 0.073 g KD/g of mouse/day.

The average dietary fluxes for individual ingredients for mice on a diet are then calculated by using the total dietary fluxes and the composition of the diets (CD: PicoLab® Rodent Diet 20 5053; KD: Bio-serv F3666 ketogenic diet). The diet formulas report combined glutamine + glutamate, which provides an upper bound on glutamine. The diet formulas specify the content of total saturated fatty acids, but not specific saturated fatty acids. Palmitic acid (C16:0) and oleic acid dietary flux is based on our measurement that palmitic acid and stearic acid (C18:0) are the main saturated fatty acids, with a ratio of 2.4:1 for CD and 1.7 for KD. The palmitic acid dietary flux was calculated based on this number for each diet. Other amino acids (other AAs) include arginine, aspartate, asparagine, cystine, proline, and tyrosine. The dietary flux for other fatty acids (other FAs) was calculated as the total dietary flux of fatty acids minus the sum of the dietary fluxes of palmitic acid, oleic acid, and linoleic acid.

### Definition of *F*_*circ*_ and *F*′_*circ*_

To measure the circulatory turnover flux of a nutrient with a carbon number of *C*, the uniformly ^13^C-labeled form of the nutrient is infused. At steady state, the fraction of the mass isotopomer [*M* + *l*] of the nutrient in serum is measured as *L*_[*M*+*l*]_. (Note that there are a total number of *C* + *1* mass isotopomer for the nutrient, such that *l* is from 0 to *C*.) The circulatory turnover flux *F*_*circ*_ is defined as

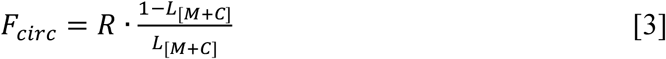

where *R* is the infusion rate of the labeled tracer.

The carbon-atom circulatory turnover flux *F*′_*circ*_ of the nutrient is defined as

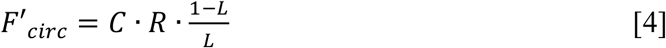

where *L* is the fraction of labeled carbon atoms in the nutrient, or mathematically

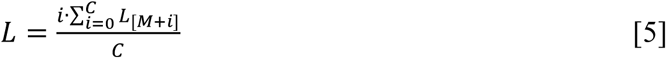

As the definitions show, *F*_*circ*_ measures the turnover of the whole carbon skeleton of the molecule while *F*′_*circ*_ measures the turnover of the carbon atom in the molecule.

### Definition of Normalized Labeling

In the infusion of a ^13^C-labeled tracer *X*, the normalized labeling of a nutrient *Y* is defined as 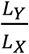, where *L*_*X*_ and *L*_*Y*_ are the fraction of labeled carbon atoms for *X* and *Y*, respectively, defined in Eqn. [5].

### Calculation of direct contributions with the constraint of non-negative values

As explained in the main text, by constructing a set of linear equations, we can solve them for the direct contribution from each of the circulating nutrients to a specific circulating nutrient (Eqn. [1] or the direct contribution from each circulating nutrient to the tissue TCA cycle (Eqn. [2]). Due however to measurement errors, the calculated direct contributions are sometimes negative, albeit with small values. To obtain the direct contributions that best reflect the biological fluxes, we use an optimization procedure to search for the set of direct contributions (represented as vector *f*) that minimize the Euclidean distance between the predicted labeling (represented as vector *M* · *f*) and the measured labeling (represented as vector *L*), under the constraint of non-negative values for *f*. Or mathematically,

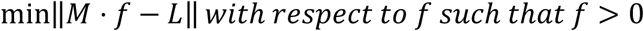

The optimization was performed using Matlab function fmincon.

To estimate the errors for the values in *f*, we used the bootstrapping procedure to randomly sample values for the matrix *M* and *L* (with n=100), which were then used for obtaining values of *f*. The standard deviation of the resulting values of each direct contribution was taken as its error.

### Quantification of Absolute Flux between Circulating Nutrients

See Suppl. Note in the SI.

## DATA AND CODE AVAILABILITY

El-Maven (Elucidata) was used for analysis of LC-MS data. Isotope labeling was corrected using AccuCor Isotope Natural Abundance Correction available on GitHub.

## SUPPLEMENTAL INFORMATION

Supplemental information includes 4 figures, 11 tables, and one supplementary note.

## ACKNOWLEDGEMENTS

S.H. is supported by the NIH grant K99DK117066. T.T. is supported by the NIH grant 1F32DK118856-01A1. C.J. is a postdoctoral fellow of the American Diabetes Association (1-17-PDF-076). W.L. is supported by the NIH grant CA211437. This work was supported by NIH Pioneer award 1DP1DK113643 and Diabetes Research Center grant P30 DK019525. We thank the University of Pennsylvania Diabetes Research Center (DRC) for the use of the Rodent Metabolic Phenotyping Core (P30-DK19525).

We thank members of the Rabinowitz lab for scientific discussions.

## AUTHOR CONTRIBUTIONS

S.H., A.J.C, and J.D.R. designed the study. S.H. performed most experiments and data analysis in mice fed on high carbohydrate diet. A.J.C. performed most experiments and data analysis in mice fed on ketogenic diet. X.Z. performed isotopic labeling measurement of acetate. L.Y. contributed to isotope tracing experiments in mice fed on ketogenic diet. X.Z., X.L., C.B., Z.Z, C.J. and L.W. contributed to isotope tracing experiments in mice fed on high carbohydrate diet. S.H. and J.D.R. performed the flux modeling. L.W. and W.L. contributed to LC-MS analysis of metabolites. A.J.C., J.R., and J.B. contributed to metabolic measurements of mice. S.H., A.J.C., and J.D.R. wrote the manuscript. All authors discussed the results and commented on the manuscript.

## SUPPLEMENTAL INFORMATION

### SUPPLEMENTAL FIGURES

**Figure S1:**
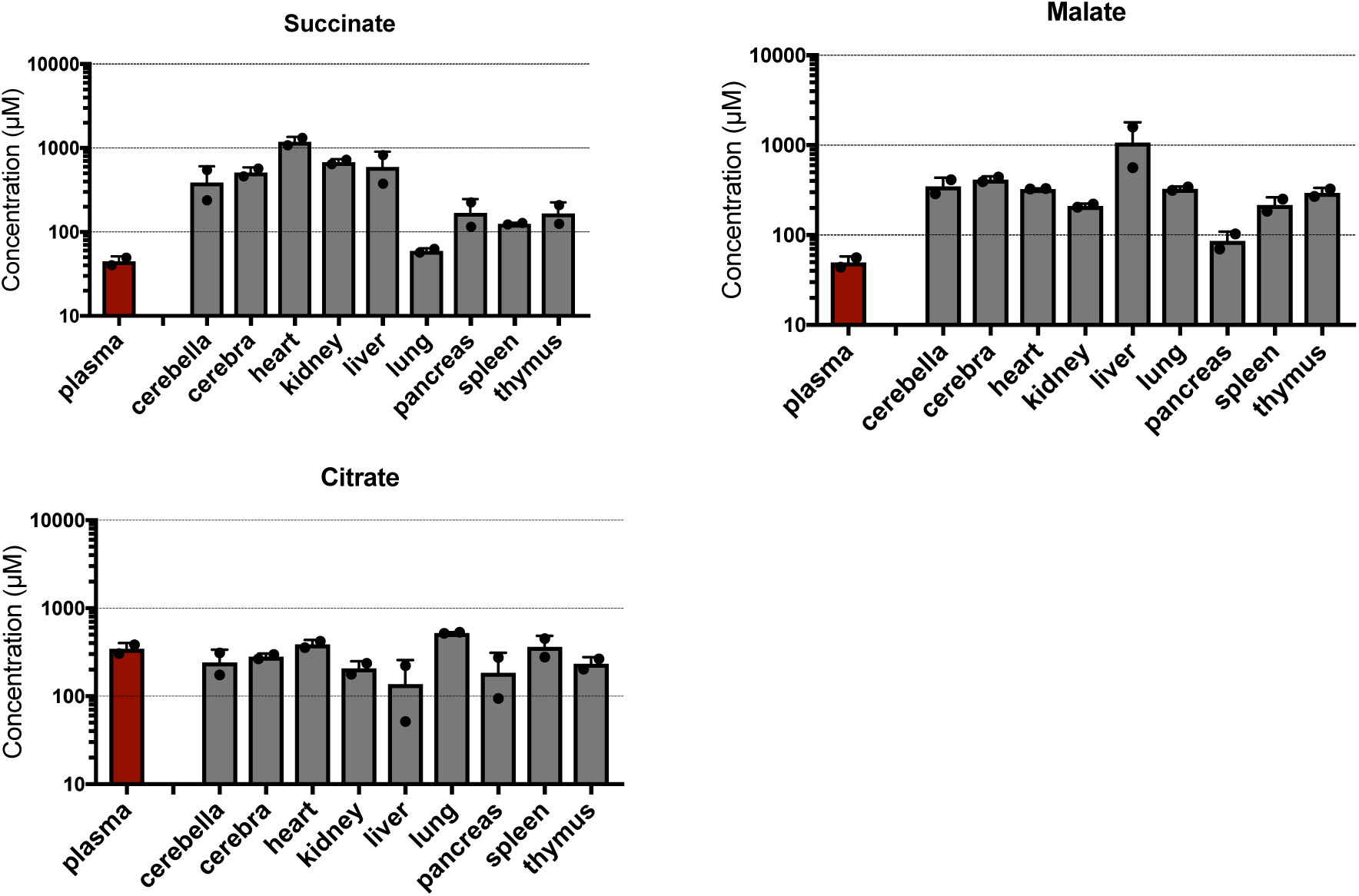
Concentrations of succinate and malate are generally higher in tissues than in plasma, while citrate has similar concentration between tissues and in plasma. Data are from the Mouse Multiple tissue Metabolome Database (http://mmdb.iab.keio.ac.jp). Values are mean ± SD (n=2 mice). Note that Y-axis is logarithmic scale. (Adapted from Hui et al. 2017).

**Figure S2:**
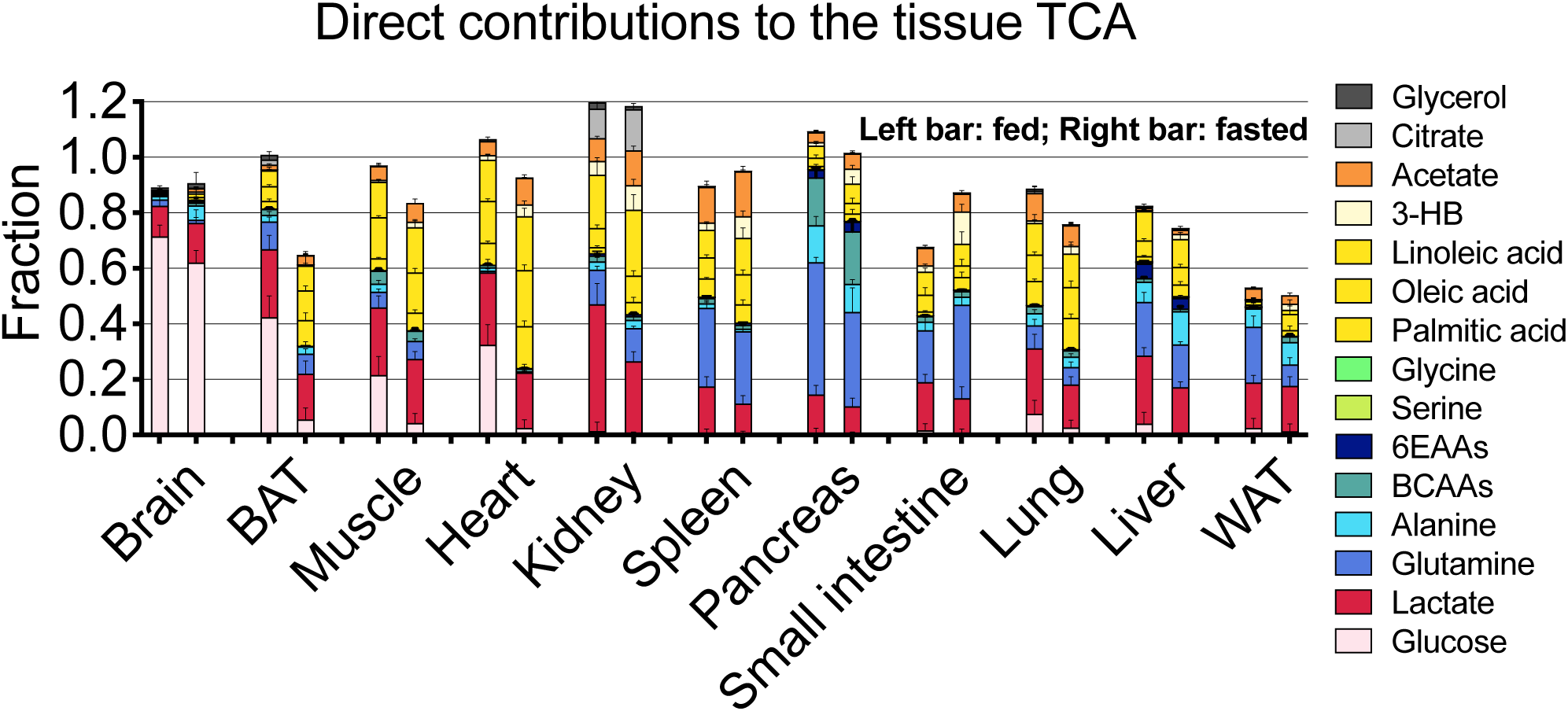
Direct TCA contributions using tissue succinate labeling measurements show same trends as those obtained using tissue malate labeling data (in Figure 4C). Data are for 8-h fasted mice and 3-h refed mice on carbohydrate diet.

**Figure S3.**
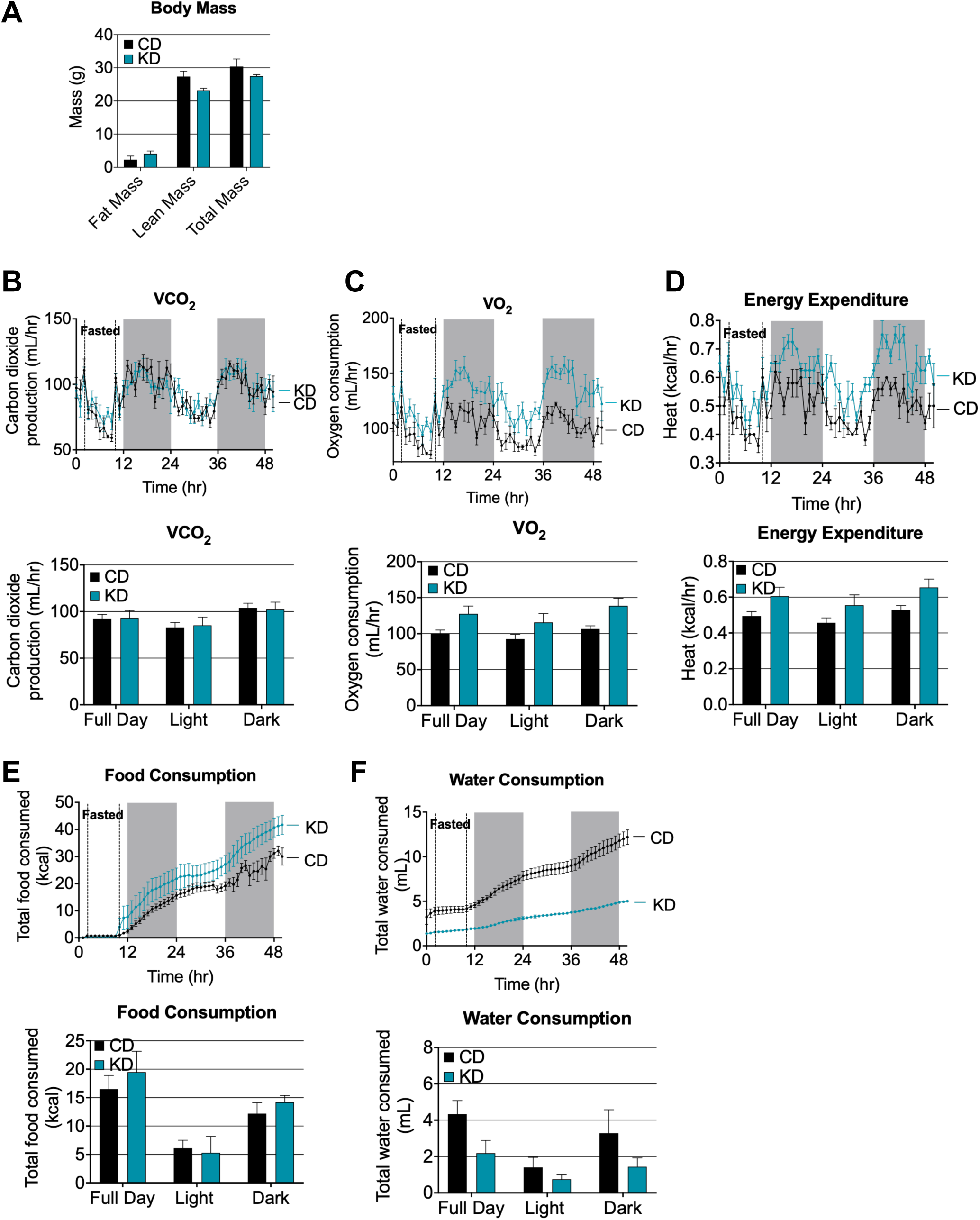
Metabolic and physical parameters of mice fed carbohydrate diet (CD; N=5) and ketogenic diet (KD; N=4). Bar graphs show mean ± SD, excluding the fasting period. (A) Body composition. (B) Oxygen consumption. Shaded areas indicate the dark periods. Mice were fasted for 8 h in the first light period. (C) Carbon dioxide consumption. (D) Energy expenditure. (E) Food consumption. (F) Water consumption.

**Figure S4.**
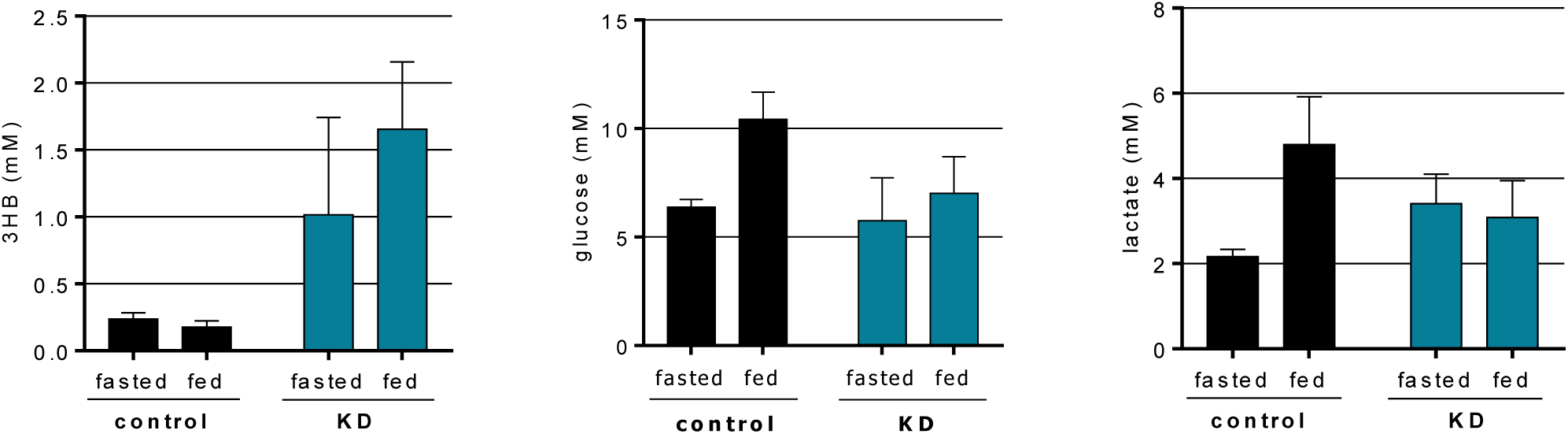
Serum concentrations of 3-hydroxybutyrate, glucose and lactate in mice fed on the carbohydrate diet (N=5) and ketogenic diet (N=4) (mean ± SD).

### SUPPLEMENTAL TABLES

**Table S1.** *F*_*circ*_, 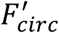, and *F*_*diet_avg*_ for fasted mice on carbohydrate diet. Related to Figure 1B, C.

| Nutrients | $F_{circ}$ | SEM | $F'_{circ}$ | SEM | N | $F_{diet\_avg}$ | |
| --- | --- | --- | --- | --- | --- | --- | --- |
|  | nmol/min/g |  | nmol carbon/min/g |  |  | nmol/min/g | nmol carbon/min/g |
| glucose | 151 | 10 | 752 | 50 | 22 | 234 | 1403 |
| lactate | 374 | 23 | 854 | 52 | 24 | 0 | 0 |
| glutamine | 46 | 2 | 219 | 10 | 5 | <24 | <118 |
| alanine | 81 | 5 | 242 | 16 | 8 | 10 | 31 |
| 3HB | 42 | 5 | 168 | 18 | 9 | 0 | 0 |
| palmitate | 24 | 1 | 383 | 23 | 7 | 2 | 33 |
| oleate | 35 | 2 | 630 | 39 | 6 | 3 | 50 |
| linoleate | 45 | 2 | 801 | 41 | 6 | 6 | 112 |
| Serine | 20 | 3 | 53 | 7 | 5 | 8 | 23 |
| Glycine | 20 | 2 | 39 | 4 | 5 | 10 | 20 |
| Citrate | 11 | 1 | 67 | 7 | 13 | 0 | 0 |
| Acetate | 56 | 8 | 112 | 15 | 7 | 0 | 0 |
| Glycerol | 67 | 4 | 201 | 11 | 8 | 5 | 16 |
| BCAAs | 31 | 2 | 177 | 11 | 7 | 22 | 126 |
| 6EAAs | 34 | 2 | 183 | 12 | 6 | 23 | 143 |
| Other AAs | 70 | 9 | 374 | 52 | 3 | 32 | 163 |
| Other FAs |  |  |  |  |  | 5 | 90 |

Table S2. Normalized labeling data for 15 tracer infusions for both fasted and fed mice on carbohydrate diet. Related to Figure 2C, 3C, and 4C.

See Table S2.xlsx for the table.

The data table includes the normalized labeling data for each of the 15 tracer infusion experiments. For each infusion, measurements are provided for 15 circulating metabolites in serum and for malate and succinate in 11 tissues. Values are mean ± SEM.

Table S3. Direct contributions to circulating nutrients for fasted mice on carbohydrate diet. Related to Figure 2C, 3C.

See Table S3.xlsx for the table.

The data table includes the direct contributions for each of the 15 circulating nutrients from the other 14. Errors are determined by bootstrapping (see Methods).

**Table S4:**
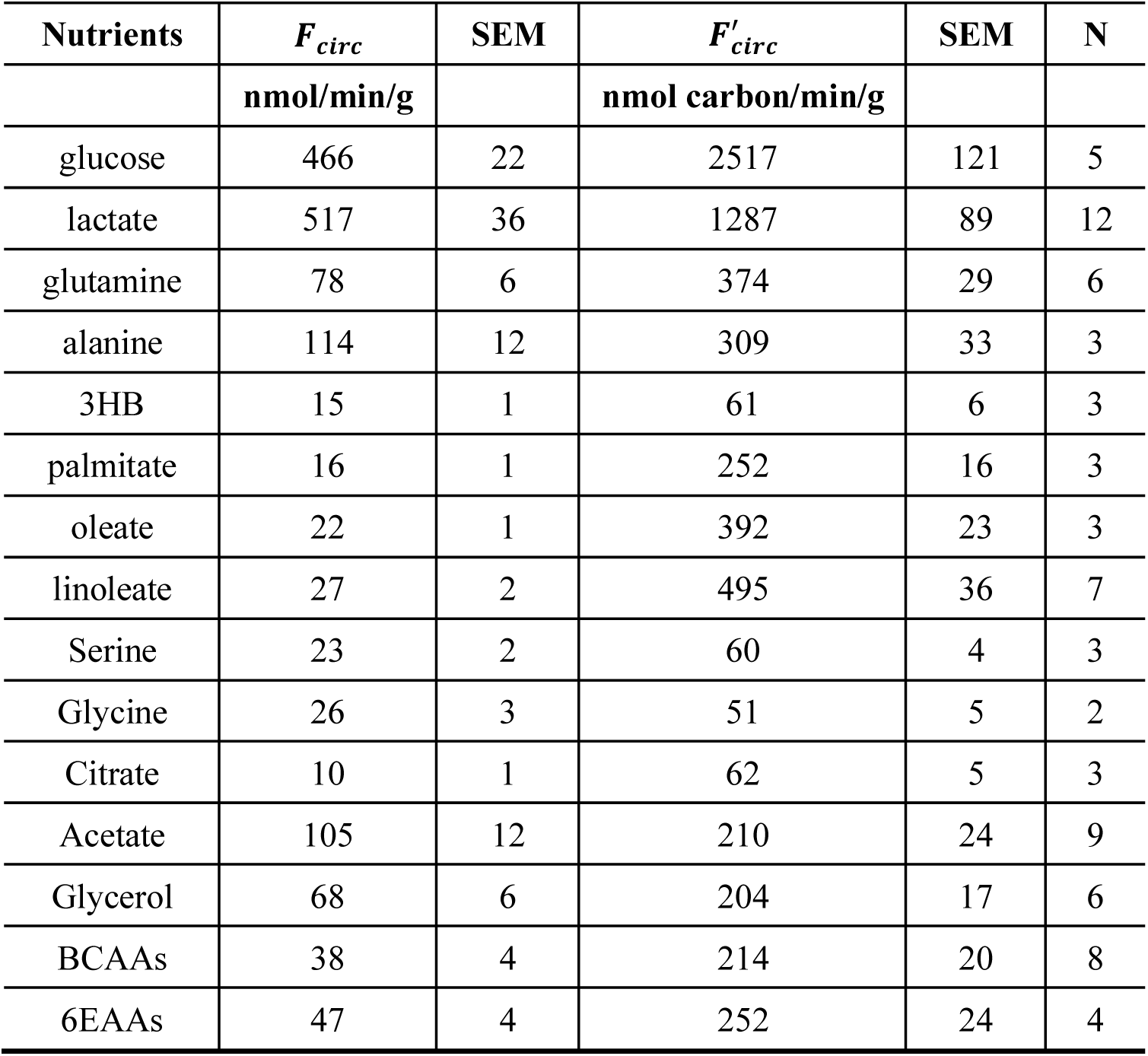
Values of *F*_*circ*_ and 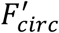, for fed mice on carbohydrate diet. Related to Figure 3B.

Table S5. Direct contributions to circulating nutrients for fed mice on carbohydrate diet. Related to Figure 3C.

See Table S5.xlsx for the table.

The data table includes the direct contributions for each of the 15 circulating nutrients from the other 14. Errors are determined by bootstrapping (see Methods).

Table S6. Direct contributions to tissue TCA cycle in both fasted and fed mice on carbohydrate diet. Related to Figure 4C.

See Table S6.xlsx for the table.

The data table includes the direct contributions to the TCA cycle of 11 tissues from 15 circulating nutrients. Errors are determined by bootstrapping (see Methods).

**Table S7.** Composition of carbohydrate and ketogenic diets.

|  | Carbohydrate diet |  | Ketogenic diet |  | Change in dietary flux |
| --- | --- | --- | --- | --- | --- |
|  | by weight | by calories | by weight | by calories |  |
| **Carbohydrates** | 50.3% | 62.1% | 3.2% | 1.8% | 25-fold less in KD |
| **Fat** | 4.8% | 13.2% | 75.1% | 93.4% | 10-fold more in KD |
| **Protein** | 20.0% | 24.7% | 8.6% | 4.7% | 4-fold less in KD |
| **Rest** | 24.9% | 0.0% | 13.1% | 0.0% |  |

To calculate the change in dietary flux, we utilize our measured the calorie intake for mice on the two diets: 0.39±0.02 Calories/day/g for mice on CD (N=5) and 0.53±0.04 Calories/day/g for mice on KD (N=4). For example, for carbohydrates, the change in dietary flux from CD to KD is given by (0.28x 62.1%)/(0.39 x 1.8%) = 25.

**Table S8.** *F*_*circ*_, 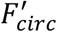, and *F*_*diet_avg*_ for both fasted and fed mice on ketogenic diet. Related to Figures 5A and 5B.

| | Fasted | | | | | $F_{diet\_avg}$ | |
| --- | --- | --- | --- | --- | --- | --- | --- |
| | $F_{circ}$ | SEM | $F'_{circ}$ | SE<br>M | N | nmol/min/g | nmol carbon/min/g |
|  | nmol/m<br>in/g |  | nmol<br>carbon/min<br>/g |  |  |  |  |
| glucose | 66 | 3 | 302 | 17 | 5 | 9 | 54 |
| lactate | 292 | 12 | 688 | 22 | 4 | 0 | 0 |
| glutamine | 49 | 6 | 242 | 27 | 4 | <6 | <30 |
| alanine | 44 | 3 | 133 | 9 | 5 | 1 | 4 |
| 3HB | 91 | 4 | 339 | 21 | 4 | 0 | 0 |
| palmitate | 17 | 2 | 272 | 31 | 3 | 38 | 602 |
| oleate | 36 | 3 | 640 | 51 | 3 | 51 | 926 |
| linoleate | 57 | 10 | 1027 | 173 | 3 | 21 | 372 |
| Serine |  |  |  |  |  | 2 | 7 |
| Glycine |  |  |  |  |  | 1 | 3 |
| Citrate |  |  |  |  |  | 0 | 0 |
| Acetate |  |  |  |  |  | 0 | 0 |
| Glycerol | 90 | 8 | 270 | 23 | 5 | 47 | 140 |
| BCAAs |  |  |  |  |  | 7 | 39 |
| 6EAAs |  |  |  |  |  | 7 | 41 |
| Other AAs |  |  |  |  |  | 8 | 45 |
| Other FAs |  |  |  |  |  | 30 | 497 |
|  | Fed |  |  |  |  |  |  |
| | $F_{circ}$ | SEM | $F'_{circ}$ | SE<br>M | N | | |
| glucose | 77 | 3 | 420 | 18 | 3 |  |  |
| lactate | 237 | 11 | 541 | 31 | 3 |  |  |
| glutamine | 61 | 14 | 302 | 71 | 3 |  |  |
| alanine | 50 | 4 | 151 | 12 | 3 |  |  |
| 3HB | 104 | 1 | 396 | 6 | 3 |  |  |
| palmitate | 22 | 1 | 351 | 15 | 3 |  |  |
| oleate | 40 | 0 | 716 | 0 | 3 |  |  |
| linoleate | 45 | 2 | 810 | 36 | 3 |  |  |
| Glycerol | 178 | 12 | 533 | 35 | 3 |  |  |

Table S9. Normalized labeling data for 9 tracer infusions for both fasted and fed mice on ketogenic diet

See Table S9.xlsx for the table.

The data table includes the normalized labeling data for each of the 9 tracer infusion experiments. For each infusion, measurements are provided for the 9 circulating metabolites in serum and for malate in 11 tissues. Mean ± SEM.

Table S10: Direct contributions to circulating nutrients for both fasted and fed mice on ketogenic diet. Related to Figure 5C.

See Table S10.xlsx for the table.

The data table includes the direct contributions for each of the 9 circulating nutrients from the other 8. Errors are determined by bootstrapping (see Methods).

Table S11: Direct contributions to tissue TCA cycle in both fasted and fed mice on ketogenic diet. Related to Figure 6B.

See Table S11.xlsx for the table.

The data table includes the direct contributions to the TCA cycle of 8 tissues from the 9 circulating nutrients. Errors are determined by bootstrapping (see Methods).

### SUPPLEMENTAL NOTE

#### Quantifying fluxes between circulating nutrients

In this note, we explain the procedure for calculating fluxes between circulating nutrients. The input data for this calculation are (i) an *n* x *n* matrix *M*, that reflects the extent to which infusion of any nutrient *i* (of *n* total nutrients of interest) labels every other circulating nutrient *j* (“the inter-labeling matrix”) (note that generation of this matrix requires *n* isotope tracer infusion experiments) and (ii) the carbon-atom circulatory turnover flux 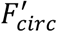 for each circulating nutrient *i* (defined as 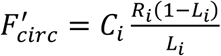 where *C*_*i*_ is the number of carbon atoms in one molecule of nutrient *i, L*_*i*_ is the fraction of labeled carbon atoms in the nutrient *i*, and *R*_*i*_ is the infusion rate of uniformly ^13^C-labeled *i*). The procedure first uses *M* to calculate the direct contributions to each nutrient *i* from all other circulating nutrients *j*, creating a new n x n matrix *N* whose entries *N*_*ij*_ reflect the (fractional) direct contributions of circulating nutrient *j* to circulating nutrient *i*. It then utilizes the matrix *N* and the 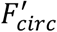 to calculate the direct contributing fluxes from any circulating nutrient to any other circulating nutrient, resulting in a complete determination of the inter-converting fluxes between circulating nutrients.

#### Calculating the direct contributions between circulating nutrients

As detailed in the main text, by constructing a set of linear equations from the inter-labeling matrix *M*, we can calculate the direct contribution to any one circulating nutrient from the other circulating nutrients. The direct contribution from nutrient *j* to nutrient *i* is defined as the fraction of *i* that comes directly from *j*, e.g., 0.7 (or 70%) of circulating lactate comes directly from circulating glucose. The results can be organized as a second *n* × *n* matrix (denoted as *N*), with the non-diagonal entry *N*_*ij*_ representing the direct contribution from nutrient *j* to nutrient *i*, or

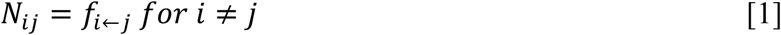

For the diagonal entries in *N*, we define

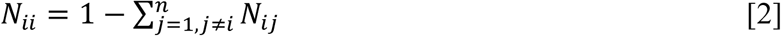

which represents the direct contribution from sources other than circulating nutrients (i.e., nutrient storages). We thus denote the diagonal entries

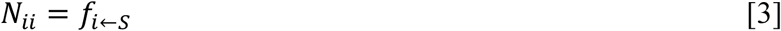

#### Calculating the direct contributing fluxes to a circulating nutrient

The matrix *N* contains the direct contribution values as fractions. To obtain the direct contribution in flux units (e.g., nmol carbon/min/g), we utilize the measured carbon-atom circulatory turnover flux values 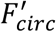. The calculation is more complex than just multiplying the direct contributions by 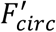, however, because only unlabeled inputs to a metabolite contribute to 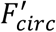. The total flux also includes metabolic cycles where an infused labeled metabolite may be regenerated in labeled form from its own products. This requires multiplying 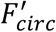 by an adjustment factor *c* ≥ 1. For illustration, we use a simple case where there are only 3 circulating nutrients (Fig. 1).

**Figure S6.**
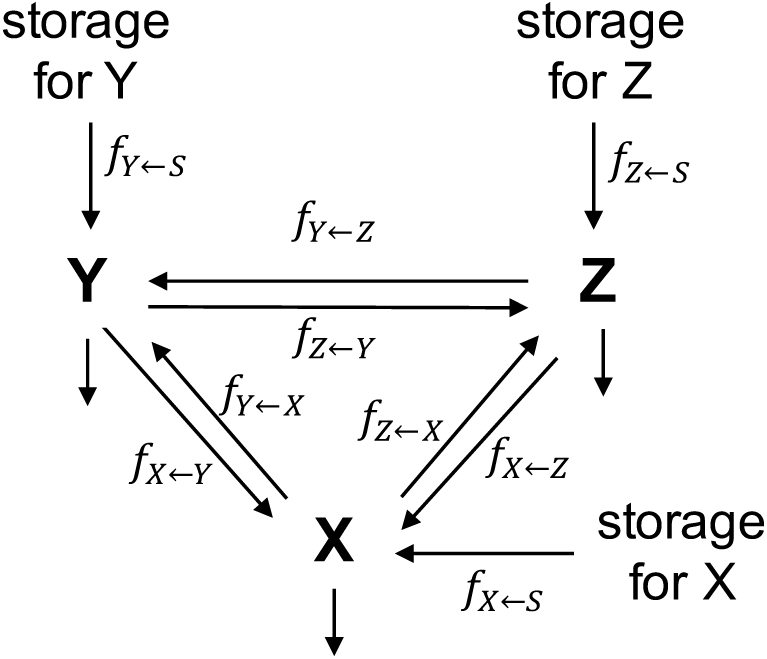
A simple case illustrating the inter-conversion between 3 circulating nutrients.

We first focus on one nutrient X and calculate the direct contributing fluxes to it: the flux from Y to X (*J*_*X*←*Y*_), the flux from Z to X (*J*_*X*←*z*_), and the flux from any unmeasured, unlabeled inputs to X (*J*_*X*←*s*_) (the most important unlabeled inputs are diet and storage polymers). We denote *J*_*X*_ as the total flux that goes through X, meaning it is equal to the sum of the incoming fluxes to X (which at steady-state is also equal to the sum of the outgoing fluxes from X). Mathematically,

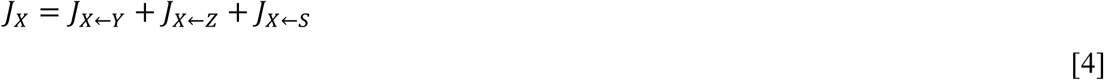

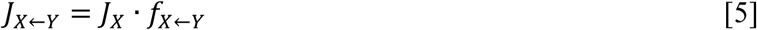

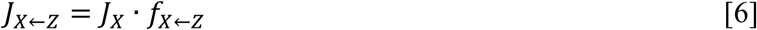

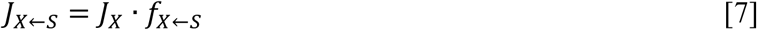

To calculate *J*_*X*_, we begin by expressing 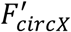 in terms of the fluxes on the network shown in Fig. 1. 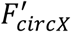 reflects flux of unlabeled carbon into X. Such unlabeled carbon can come from multiple sources.

Path 1: external input to X (e.g. storage polymer, food)

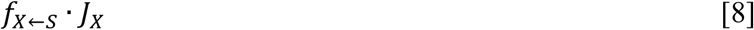

Path 2: external input to Y → X

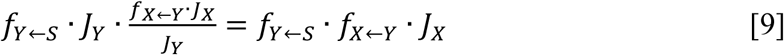

where *J*_*Y*_ is the total flux through the nutrient Y.

Path 3: external input to Y → Z → X

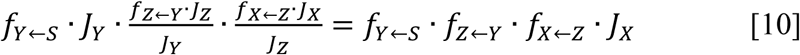

where *J*_*Z*_ is the total flux through the nutrient Z.

Path 4: external input to Z → X

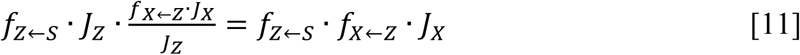

Path 5: external input to Z → Y → X

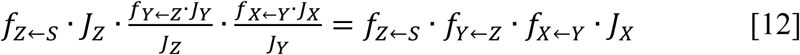

Together the 5 fluxes above give rise to 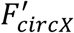, or

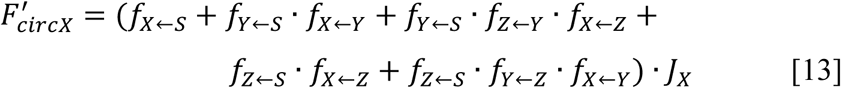

Denoting the coefficient between 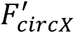 and *J*_*X*_ as *c*_*X*_, we rewrite Eqn. [13] as

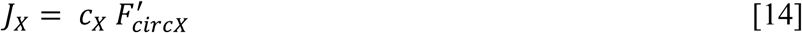

where

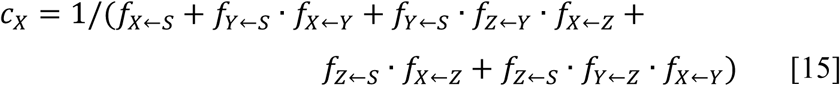

Thus, with Eqns. [14-15] and the direct contribution matrix *N*, we can calculate *J*_*X*_ for a network of 3 metabolites. Note that this equation can be generalized to any number of metabolites. Eqns. [5-7] then give the individual direct contributing fluxes to X.

